# *Rpv2* is part of a cluster of NLRs specific to *Vitis rotundifolia* and confers extreme resistance to grapevine downy mildew

**DOI:** 10.1101/2025.03.25.645223

**Authors:** Laurie Marsan, Emilce Prado, Sabine Wiedemann-Merdinoglu, Laure Schmidlin, Sophie Blanc, Marion Delame, Sylvain Schnee, Burak Arti, Guillaume Barnabé, Amandine Velt, Vincent Dumas, Didier Merdinoglu, Camille Rustenholz, Pere Mestre

## Abstract

Downy mildew caused by the oomycete *Plasmopara viticola* is one of the most important diseases affecting grapevine. Resistant varieties are an environmentally-friendly tool to control grapevine downy mildew. Efficient breeding for durable resistance requires knowledge of the underlying mechanisms. Here we aimed at identifying the molecular basis of *Rpv2*, a gene for extreme resistance to downy mildew derived from *Vitis rotundifolia*, and at characterizing its effect on pathogen development. Individuals from two populations segregating for *Rpv2* were evaluated for resistance to downy mildew and genotyped. Following genetic mapping, markers flanking *Rpv2* were used to screen new populations and identify recombinant individuals. Sequencing of recombinants and *in silico* chromosome painting was used to reduce the interval containing *Rpv2*. Comparative genomics inside the *Vitaceae*, involving *de novo* assembly of the *V. rotundifolia* Regale genome, allowed narrowing-down the list of candidate genes. We restrict *Rpv2* to a 250 kb genomic region that contains two resistance genes of the NLR type. Comparative genomics analyses could not find orthologs of both NLRs in the other *Vitis* species studied. We also show that *Rpv2*-mediated resistance leads to pathogen arrest early in the infection cycle. Our results show that *Rpv2* belongs to the NLR family of resistance genes, contributing thus to understand the potential and risks of its use in breeding programs and suggesting that combining NLR-type genes may lead to durable resistance

**KEY MESSAGE:** The *Rpv2* locus for extreme resistance to grapevine downy mildew is mapped to a 250 kb genomic region containing two NLR-type genes specific to *V. rotundifolia*.

## INTRODUCTION

Grapevine is a culture of economic importance cultivated in temperate or warm regions all around the world for the production of wine, fresh fruit, raisins and juice. Other than its economic importance, grapevine cultivation and wine production are deeply rooted in human culture since ancient times, and still today the socio-cultural aspect of wine is patent, its consumption being often associated with social interaction and celebration.

Downy mildew caused by the oomycete *Plasmopara viticola* (Berk. & Curt.) Berl.& de Toni (Dick 2002) is one of the major threats to grapevine. *P. viticola* was introduced in France from North America during the 19th century and rapidly spread across Europe and, from there, to all other grapevine-growing areas of the world (Gessler et al. 2011; Fontaine et al. 2021). *P. viticola* is an obligate biotroph that can infect all green tissues of the grapevine, particularly leaves, reducing both the functional green leaf area and the assimilation rate of the remaining green leaf area (Moriondo et al. 2005). Inflorescences and young bunches can also be infected by downy mildew, leading to significant losses of productivity and quality (Kennelly et al. 2005).

Most traditional cultivated grapevine varieties belong to the *Vitis vinifera* species and are susceptible to *P. viticola*. In consequence, control of downy mildew is currently achieved mostly by regular applications of fungicides. However, the intensive use of chemicals is being progressively restricted because of their risk for human health (Inserm 2021) and their detrimental impact on the environment. Furthermore, the emergence of fungicide-resistant *P. viticola* strains decreases the efficiency of the treatments (Gisi et al. 2007; Massi et al. 2021).

Using downy-mildew resistant varieties is a cost-effective strategy harmless to the environment and human health. Creating grapevine resistant varieties implies introducing the resistance from related resistant *Vitis* species through breeding programs. Many sources of resistance to downy mildew were described in the species related to *Vitis vinifera* (Boubals 1959; Dai et al. 1995; Staudt and Kassemeyer 1995; Brown et al. 1999; Kortekamp and Zyprian 2003; Cadle-

Davidson 2008) and some of them have already been successfully introduced in grapevine to create resistant varieties (Csizmazia and Bereznai 1968; Eibach and Topfer 2003; Cadle-Davidson 2008; Merdinoglu et al. 2018). The beginning of the 21st century witnessed a progress in the study of the genetic determinism of resistance to grapevine downy mildew. To date, thirty downy mildew resistance factors (*Rpv*, for **R**esistance to ***P****lasmopara **v**iticola*) have been identified (Merdinoglu et al. 2018; Röckel 2024) but several of them confer a resistance that is too weak or only expressed in a particular genetic context. In consequence, despite the large number of sources and factors of resistance to downy mildew identified, the number of genes actually used in breeding programs is very limited. For example, varieties derived from European breeding programs rely solely on *Rpv1* (Merdinoglu et al. 2003), *Rpv3* (Bellin et al. 2009; Di Gaspero et al. 2012), *Rpv10* (Schwander et al. 2012) and *Rpv12* (Venuti et al. 2013).

The durability of resistance, which can be defined as the length of time the resistance remains effective (Johnson 1984), is an important criterion to consider, in particular for species as grapevine, where plants are meant to stay in the field for decades. Knowledge on the molecular basis of resistance makes possible understanding the mechanisms underlying plant-pathogen interactions and evaluating the risks of resistance-breaking by the pathogens. *Rpv1* and *Rpv3.1* have been identified as belonging to the NBS-LRR family (NLRs) of resistance genes (Feechan et al. 2013; Foria et al. 2020), and the other *Rpv* genes used in breeding programs are located in regions of the genome rich in NLRs. NLRs trigger defense responses following the recognition of pathogen proteins and they are renowned for being easily overcome by pathogens. Accordingly, strains of *P. viticola* have been reported to overcome the resistance conferred by Rpv3.1, Rpv10 and Rpv12 (Peressotti et al. 2010; Delmotte et al. 2014; Wingerter et al. 2021; Paineau et al. 2022). Furthermore, it has been shown that *P. viticola* Rpv3.1 resistance-breaking strains present deletions of candidate effector proteins and that those proteins are recognized by Rpv3.1 (Paineau et al. 2024). In this context of frailty of resistance genes, identifying and characterizing new resistance genes usable in breeding programs is thus crucial not only to diversify the resistance factors but also to choose the combination of genes that would lead to a more efficient and durable resistance through a strategy of pyramiding.

*Vitis rotundifolia* is a species belonging to the *Muscadinia* sub-genus and displaying 2n=40 chromosomes instead of 2n=38 chromosomes for the *Euvitis* sub-genus. It is native from the southern United States of America, where it is cultivated mainly for the production of fresh fruit, juice and jelly. *V. rotundifolia* is described as resistant to several grapevine pests and diseases, like nematodes and downy and powdery mildews (Boubals 1959; Olmo 1971; Staudt and Kassemeyer 1995; Diez-Navajas et al. 2008; Riaz et al. 2011; Blanc et al. 2012). Resistance to downy mildew is total, i.e., there is no pathogen sporulation or plant cell death following inoculation. Genetic analysis of the resistance to downy mildew in a population derived from *V. rotundifolia* cv Dearing led to the identification of *Rpv1*, located in chromosome 12 (Merdinoglu et al. 2003), but this gene confers partial resistance and thus does not explain on its own the total resistance phenotype. Previous work from our laboratory using a different population derived from *V. rotundifolia* cv Trayshed resulted in the identification of a second locus in chromosome 18 with a strong effect, which was named *Rpv2* (Wiedemann-Merdinoglu et al. 2006). *Rpv2* is located in the same region of chromosome 18 as *Run2.2*, a gene for resistance to powdery mildew from *V. rotundifolia* cv Trayshed (Riaz et al. 2011); with our current knowledge it remains possible that resistance to both pathogens is actually conferred by the same gene. Besides, the phenotype of strong resistance conferred by *Rpv2*, different from that of other *Rpv* genes, raises the possibility that *Rpv2* is different to an NLR. To shed light on these questions, here we describe the genetic mapping of *Rpv2* and the characterization of its effect on *P. viticola* infection. Also, in order to clarify the relative positions of *Rpv2* and *Run2.2*, we phenotype an advanced breeding population for resistance to downy and powdery mildews.

Finally, by performing fine genetic mapping and using newly created and available genomic resources, we clarify the molecular basis of the *Rpv2*-mediated resistance.

## MATERIALS AND METHODS

### Plant material

The *Vitis rotundifolia*-derived pseudo-BC1 and pseudo-BC4 populations have been described by (Delame et al. 2019). Briefly, the pseudo-BC1 population was derived from a cross between a hybrid named 8624 (*M. rotundifolia cv.* Trayshed x (*V. vinifera cv* Carignan x *V. vinifera cv.* Cabernet Sauvignon)) as female parent and *V. vinifera cv.* Cabernet Sauvignon as pollen donor. The pseudo-BC4 population was derived from the cross between the Rpv2-containing BC3 individual 1771P as female parent and *V. vinifera cv* Nebbiolo as male parent.

Two pseudo-BC5 populations were obtained by crossing the *Rpv2*-containing BC4 individuals 1128R and 1141R as female parents with, respectively, *V. vinifera* cv Semillon and cv Muscat d’Alexandrie as pollen donors. The populations comprised 24 and 83 individuals, respectively.

Selfing populations were obtained for the BC3 individual 1771P and the *Rpv2*-containing BC4 individual 1149R. The populations comprised 828 and 679 individuals, respectively. The pedigrees of the populations used in this study are shown in Fig. S1.

Seedlings were grown in the greenhouse either on rockwool blocks (D4 40/35*1 MID S DEL WG (8H), R Grodan, Roermond, The Netherlands) stalked on 3 m nylon wires, or on peat (RHP25, Klasmann-Deilmann France Sarl) and then transferred to 4 L pots containing a substrate composed of 1/3 perlite and 2/3 sand. Plants were watered daily with a complete nutritive solution (5% Plant Prod 17-10-20, Fertil SAS, France; 5% Plant Prod 20–20-20, Fertil SAS, France; 1.3% Yara TeraR, KRISTA MAG, Yara, France). Biological replicates of the pseudo-BC1 plants and of susceptible and resistant controls were produced by vegetative propagation as own-rooted green cuttings maintained in pots.

### Evaluation of resistance to downy mildew

A strain of *P. viticola* collected from a susceptible *V. vinifera* in an experimental vineyard at INRAE-Colmar (France) was maintained on one-month old seedlings of *V. vinifera cv.* Muscat Ottonel grown in the greenhouse. Sporangia were recovered from infected leaves six days after inoculation by immersion in water and gentle shaking. The concentration of the *P. viticola* suspension was measured using a cell-counting chamber.

For the pseudo-BC1 population, sixteen leaf discs of 10 mm of diameter were sampled from the fifth and the sixth expanded leaves from the apex of the grape shoots at 10-leaf stage, and placed on wet filter paper in Petri dishes the abaxial side up. For the other populations, three 20 mm leaf discs were excised from the fifth leaf and placed on twelve-well plates, on wet paper discs loaded on agar solution (10 g/L), abaxial side up. The leaf discs were artificially inoculated by spraying with a *P. viticola* suspension at 10^5^ sporangia/ml. Twelve-well plates and Petri dishes were then sealed and incubated in a growth chamber at 21°C and a photoperiod of 18 h light/6 h darkness. Scoring of each inoculated leaf disc was performed six days post-inoculation (dpi). Individuals from the pseudo-BC1 population were scored for the parameters described in Table S1, aiming at measuring the general level of resistance (OIV452), the effect of resistance on the sporulation of *P. viticola* (S, SQ, DS, NBSCP) and the necrotic symptoms produced in response to infection (N, NSTO). Individuals from the pseudo-BC4 population were scored for the parameters described in Table S2. For each plant of the pseudo-BC1 population, the number of sporangia produced on leaf discs (NBSCP) was measured with a Z2 Coulter Cell Counter (Beckman Coulter) by sampling 10 leaf discs in 10 mL solution, whereas for the pseudo-BC4 population, the number of sporangia produced per on each leaf disc (SPNB) was measured with a Scepter ^TM^ 2.0 Cell Counter (Merck Millipore) by adding 1 mL to each leaf disc present in an Eppendorf tube. Sporulating area (SA) was assessed by image analysis (Peressotti et al. 2011).

Two independent inoculations were performed for each population and the average score between repeats was used for mapping purposes.

Evaluation of resistance to powdery mildew was performed by measuring the general level of resistance (OIV455) following natural infection in the greenhouse.

### Cytological observations

Leaf discs were stained using the Blankophor and KOH-aniline blue methods as previously described (Diez-Navajas et al. 2007). Following staining, leaf discs were observed with a Leitz Weztlar epifluorescence microscope (excitation filter 340-380 nm; dicroic mirror 400 nm; barrier filter 430 nm). Blankophor staining was used to quantify spore encystement and number of stomata, while KOH-aniline blue staining was used to observe intercellular (intra-tissular) infection structures. Three independent leaf discs were used per genotype. For each leaf disc, stomata, encysted spores and pathogen structures corresponding to different pathogen developmental stages (MYC0 to MYC4) were counted in five different ocular fields.

### Simple sequence repeat (SSR) marker analysis

Genomic DNA was extracted from approximately 80 mg of young expanding leaves collected in the greenhouse using the Qiagen DNeasy® Plant Mini Kit (Qiagen S.A., Courtaboeuf, France) as described by the supplier. All microsatellite loci were amplified in a 8 µl-vol reaction mixture containing 2.5 mM MgCl2, 150 µM dNTPs, 0.25 µM of the fluorescent-labelled primer (FAM, HEX or NED), 0.5 µM of the unlabelled primer, 0.025 U/µl of AmpliTaq Gold DNA polymerase, 1 ng/µl of grapevine DNA, 1x Gold Buffer. Amplifications were performed on a Perkin Elmer 9700 thermocycler programmed as follows: 10 min at 94 °C followed by 35 cycles of 45 s at 92 °C, 60 s at 57 °C, 90 s at 72 °C and a final step of 5 min at 72 °C. Up to three different primer pairs were mixed in the same PCR reaction taking into account the size of the amplified fragments and/or the labelling of the primers (Merdinoglu et al. 2005). The mix of four PCR products, according to their size and labelling, allowed us to analyse of up to twelve markers in one injection. After a 1/5 dilution in water, 1 µl of the PCR products was added to a 19 µl mixture of formamide and HD400-ROX as internal size standard. The mix was then denatured for 3 min at 92 °C. All products for amplification and electrophoresis were obtained from Applied Biosystems. Microsatellite fragments were resolved on an automated ABI Prism 310 Genetic Analyser (Applied Biosystems) using a 36 cm capillary filled with the POP-4 polymer. Electrophoregrams were analysed using Genescan^TM^ 3.1 (Applied Biosystems). Alleles were identified using Genotyper^TM^ 2.5.2 (Applied Biosystems) and their size determined using the HD400-ROX internal size standard.

### Genetic mapping

Three hundred and twenty-eight primer pairs flanking microsatellite loci from marker sets VVS (Thomas and Scott 1993), VVMD (Bowers et al. 1996; Bowers et al. 1999), VrZAG (Sefc et al. 1999), VMC (Vitis Microsatellite Consortium, coordinated by Agrogene, Moissy Cramayel, France), UDV (Di Gaspero et al. 2005) and VVI (Merdinoglu et al. 2005) were screened for informative segregation on parents and six randomly chosen individuals from each population. One-hundred and thirty-two and 138 polymorphic SSR markers were used to obtain the genetic maps for the pseudo-BC1 and pseudo-BC4 populations respectively.

To develop new microsatellite markers in the *Rpv2* region on LG 18, the genome sequence of PN40024 12X version (https://urgi.versailles.inra.fr/Species/Vitis/Data-Sequences/Genome-sequences) in the region between SSR markers VVMD17 and VVIn16 was scanned for microsatellite sequences using WebSat (Martins et al. 2009)(https://bioinfo.inf.ufg.br/websat/) and primers were developed using Primer3 (Rozen and Skaletsky 2000). Primer sequences are shown in Table S3.

Segregation patterns were assigned to each marker (<abxcd>, <efxeg>, <lmxll>, <nnxnp>, <hkxhk>) and genotypes were encoded following JoinMap 3.0 data entry notation (Van Ooijen and Voorrips 2001). Linkage analysis was performed with JoinMap 3.0, allowing the analysis of cross-pollinated populations derived from heterozygous parents and the construction of consensus linkage maps. Recombination fractions were converted into centiMorgans (cM) using the Kosambi function (Kosambi 1944). The threshold value of logarithm of odd (LOD) score was set at 4.0 to claim linkage between markers with a maximum fraction of recombination at 0.45. The goodness-of-fit between observed and expected Mendelian ratios was analyzed for each marker locus using a χ2-test. Markers showing segregation distortion were included in the final map if their presence did not alter surrounding marker order on the linkage group. Linkage groups were numbered according to the internationally acknowledged reference map (Doligez et al. 2006; Di Gaspero et al. 2007).

### QTL analysis

QTL analysis was carried out by both non-parametric Kruskal–Wallis analysis and interval mapping using the MapQTL 5.0 software (Van Ooijen 2004). The significant LOD threshold for QTL detection at p=0.05 for each linkage group was determined by 1000 permutations of the phenotypic data. Maximum LOD values were used to estimate QTL positions with a one-LOD support interval. The total phenotypic variance and the genetic variance were evaluated with analysis of variance on each progeny

### Rpv2 haplotype identification and *in silico* chromosome painting

Genomic DNA for short-read sequencing was extracted from individuals listed in Table S3 as described above. Preparation of the libraries and sequencing in pair-end 2*150nt at a minimum depth of 12X was outsourced to INRAE-EPGV.

Short reads from the five control genotypes (parents and non-recombinant genotypes) were also aligned on *V.rotundifolia* cv. Trayshed haplotype 1 and 2 to identify the haplotype that is present in the studied populations. Short reads from all samples were aligned on *V.rotundifolia* cv. Trayshed haplotype 1 for chromosome painting, *i.e.* identification of recombination breakpoints. Alignments were performed using bwa-mem2 mem command (Vasimuddin et al. 2019) and formatted using SAMtools view and sort commands (Danecek et al. 2021). Polymorphism detection as performed using GATK MarkDuplicates, HaplotypeCaller, GenomicsDBImport, GenotypeGVCFs, VariantFiltration and SelectVariants successively (van der Auwera and O’Connor 2020). Visualization of the alignments was performed on Integrative Genomics Viewer (IGV) v.2.7.2 (Robinson et al. 2011).

### *V. rotundifolia* cv Regale genome assembly

The accession of *V. rotundifolia* Regale with plant code MU.RO.6624.Col.1, was used for extraction of high molecular-weight genomic DNA. The extraction was performed by the CNRGV (Centre National de Ressources Génomiques Végétales) – INRAE Toulouse using the Genomic-tip 100/G kit (Qiagen). The SMRT library preparation and sequencing on PacBio RSII platform (P6-C4 chemistry) was done by the IGM Genomics Center at the University of California, San Diego, following the standard PacBio protocols. A total of 68X coverage was obtained in order to perform *de novo* genome assembly. Assembly polishing and finishing was performed as described by (Badouin et al. 2020) using PacBio reads and Illumina reads (SRR6729330 and SRR6729331).

### Transcriptome assembly, Regale gene annotation and expression quantification with Regale RNA-Seq reads

Gene expression in roots, young leaves, infected young leaves, floral bud, young berries and mature berries was assessed using RNA-seq. Total RNAs were extracted according to the procedure described by (Maillot et al. 2021). The RNA-seq library preparation was performed with the TruSeq Stranded mRNA Library Prep Kit (Illumina) and was sequenced in paired-end (2 × 100 bp) on HiSeq4000 platform (Illumina technology). Library preparation and sequencing was performed by the GenomEast platform—Strasbourg. A total of 1,409,781,770 reads were obtained for all samples together, used to assemble Regale transcriptome and perform gene annotation. Transcriptome was assembled using DRAP (Cabau et al. 2017) and the output fpkm1 was used for gene annotation using EuGene as described by (Badouin et al. 2020).

*V. rotundifolia* cv. Regale RNA-seq reads were aligned onto *V. rotundifolia* cv. Trayshed haplotype 1 using Star toolkit (Dobin et al. 2013). Automatic gene annotation of *V. rotundifolia* cv. Trayshed from (Cochetel et al. 2021) was inserted in Apollo (Dunn et al. 2019) and manually corrected using Regale RNA-seq alignment. Mapped reads were counted using featurecounts program (Liao et al. 2014). Gene expression was quantified with RPKM (Read per millions mapped reads) score using the transcriptomic data from Regale.

### Identification of homologous loci and genes in *Vitis vinifera* and *Vitis rotundifolia* genomes

Homologous loci identification was performed on various available genomes: Cabernet sauvignon (Massonnet et al. 2020), Nebbiolo (Maestri et al. 2022), Malbec (Calderón et al. 2024) and PN40024 (Shi et al. 2023) for *Vitis vinifera* genotypes; nine North American *Vitis* genotypes (Cochetel et al. 2023), Trayshed (Cochetel et al. 2021; Minio et al. 2022) as a control and Carlos (Huff et al. 2023) for *Vitis rotundifolia*. 1kb DNA sequence upstream and downstream of each boundary of *Rpv2* locus from Trayshed haplotype 1 sequence were used for alignment using BLAST. Each best hit was verified using reciprocal best hit method. Homologous gene identification was performed using Exonerate protein2genome command (Slater and Birney, 2005) on homologous loci. Manually annotated protein sequences from *V. rotundifolia* cv. Trayshed were used as input and a threshold of 90% identity. The longest alignments with the highest identity percentage were retained at each site. Synteny figure was drawn using gggenome tool (https://github.com/thackl/gggenomes).

For *V. rotundifolia* cv. Regale, alignment of Trayshed *Rpv2* locus was performed on Regale whole genome assembly to determine all contigs overlapping with *Rpv2* locus using nucmer (Marçais et al. 2018).

All raw sequencing data are available under the ENA EBI project PRJEB77732. Regale genome assembly and gene annotation are available at: https://entrepot.recherche.data.gouv.fr/dataset.xhtml?persistentId=doi:10.57745/Z2YMCH

## RESULTS

### Genetic mapping of extreme resistance to downy mildew

To unveil the genetic basis of the extreme resistance to downy mildew present in *V. rotundifolia*, we generated a segregating population using as resistant parent a pseudo-F1 individual, named 8624, derived from *V. rotundifolia* variety Trayshed and displaying total resistance to downy mildew. We produced a pseudo-BC1 population consisting of 177 individuals from a cross between 8624 and the susceptible parent *Vitis vinifera* cv Cabernet sauvignon.

Resistance to downy mildew in the pseudo-BC1 population was assessed following artificial inoculation of leaf discs in controlled conditions. Leaf discs were scored for different traits, including pathogen sporulation, presence of cell-death and by the OIV452 descriptor (Table S1). Several individuals showed total resistance to downy mildew (absence of sporulation). The distribution of resistance classes in the population suggested the presence of more than one locus involved in the resistance phenotype (Fig. S2).

The population was then genotyped using SSR markers. Out of 328 markers tested, 149 were heterozygous in at least one of the parents, 70 of which were heterozygous in both parents. One hundred and thirty-one markers could be mapped and assembled into 19 linkage groups. The resulting map covers 76% of the estimated 1166.8 cM reference integrated map distance and the average distance between markers is 8.58 cM (Fig. S3). Parental genetic maps were also constructed. The maternal 8624 map covers 716.7 cM with an average distance between markers of 5.4 cM and the Cabernet sauvignon map spans 946.3 cM with an average distance of 7.1 cM (Fig. S3). Marker order is consistent with their physical order in the PN40024 12X reference genome.

QTL analysis revealed two strong QTLs for several resistance traits (Fig. 1, Table S4). A QTL located on linkage group 12, at a LOD of 4.32 and explaining 12.6% of the phenotypic variation for OIV452; and a second, stronger QTL, located on linkage group 18 at a LOD of 13.98 and explaining 36.4% of the phenotypic variation for OIV452. Remarkably, the QTL on linkage group 18 had a strong effect on the presence of sporulation, at a LOD of 22.73 and explaining 55.6% of the phenotypic variance. Other minor QTLs were detected on linkage groups 7, 13 and 15 (Table S4). The QTL at linkage group 12 located at the same position as *Rpv1*, suggesting it may be the same gene or an allele. The QTL at linkage group 18 has the features of a major gene that we named *Rpv2*.

**Figure 1.**
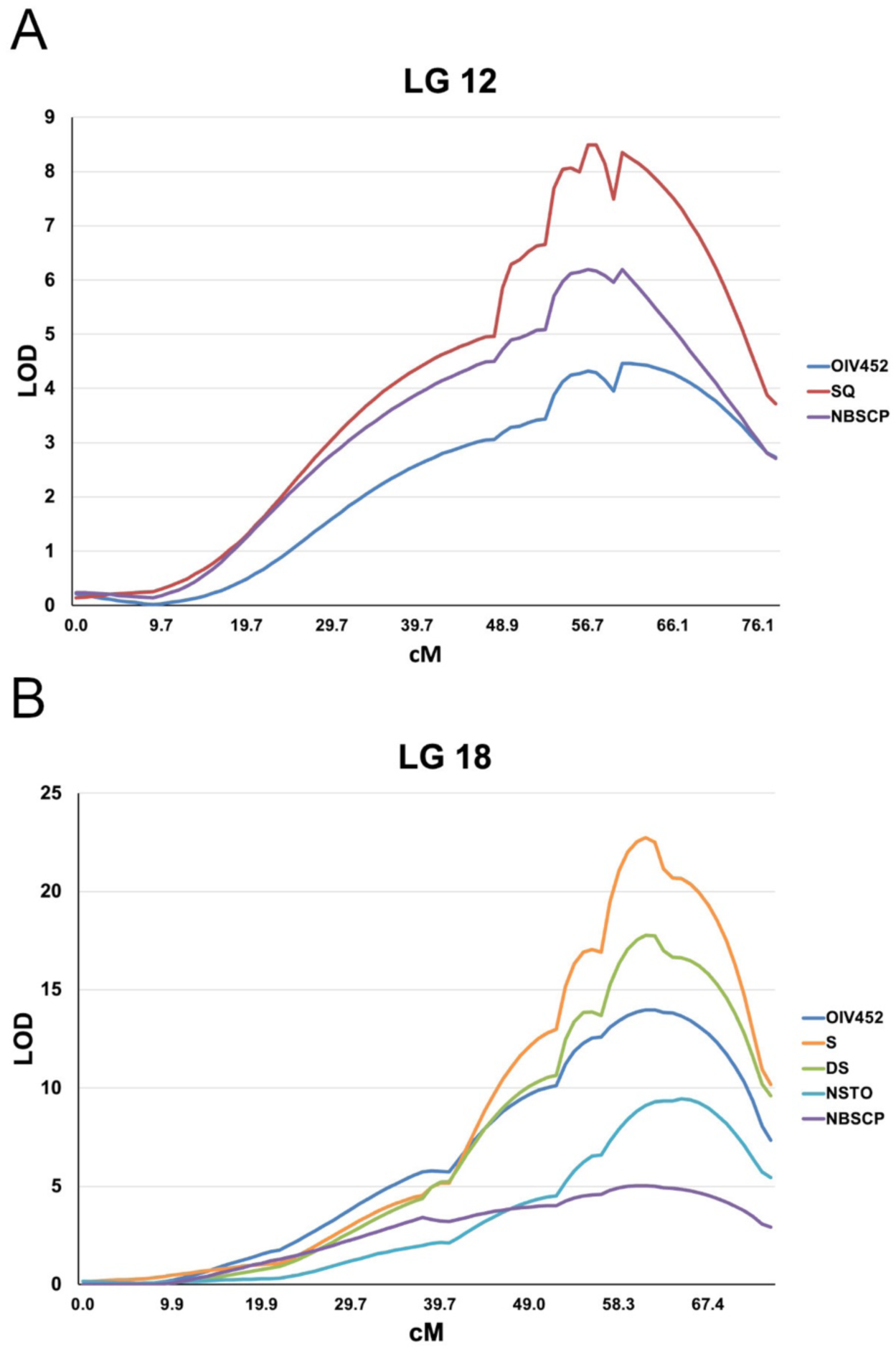
QTLs for resistance to downy mildew in the pseudo-BC1 population. Main QTLs found in linkage groups 12 (A) and 18 (B) for different variables used to evaluate resistance to downy mildew. QTL parameters, as well as other minor QTL found in the population are listed in Table S5. OIV452: symptom-based semi-quantitative score, S: sporulation; SQ: sporulation intensity; DS: percentage of sporulating leaf discs; NSTO: presence of stomatic necrosis; NBSCP: sporangia per mL.. Details about variables and scoring of are presented in Table S1.

### Effect of *Rpv2* on pathogen infection

The fact that *Rpv1* has been described as providing partial resistance to downy mildew, together with the strong effect of *Rpv2* on the presence of sporulation, led us to hypothesize that *Rpv2* alone was responsible for the absence of sporulation and the total resistance phenotype. Classifying the individuals of the BC1 population based on their genotype at the markers surrounding the *Rpv1* and *Rpv2* loci allowed to observe the effect of each individual locus on pathogen development. Plants carrying only *Rpv1* showed reduced sporulation associated to cell death (Fig. 2A), in agreement with the phenotype previously described for this gene. Plants carrying only *Rpv2* showed absence of sporulation that in some cases was associated with small necrotic spots and/or stomatal necrosis (death of the stomate companion and surrounding cells) (Fig. 2A). These results suggest that *Rpv2* confers total resistance to *P. viticola*.

**Figure 2.**
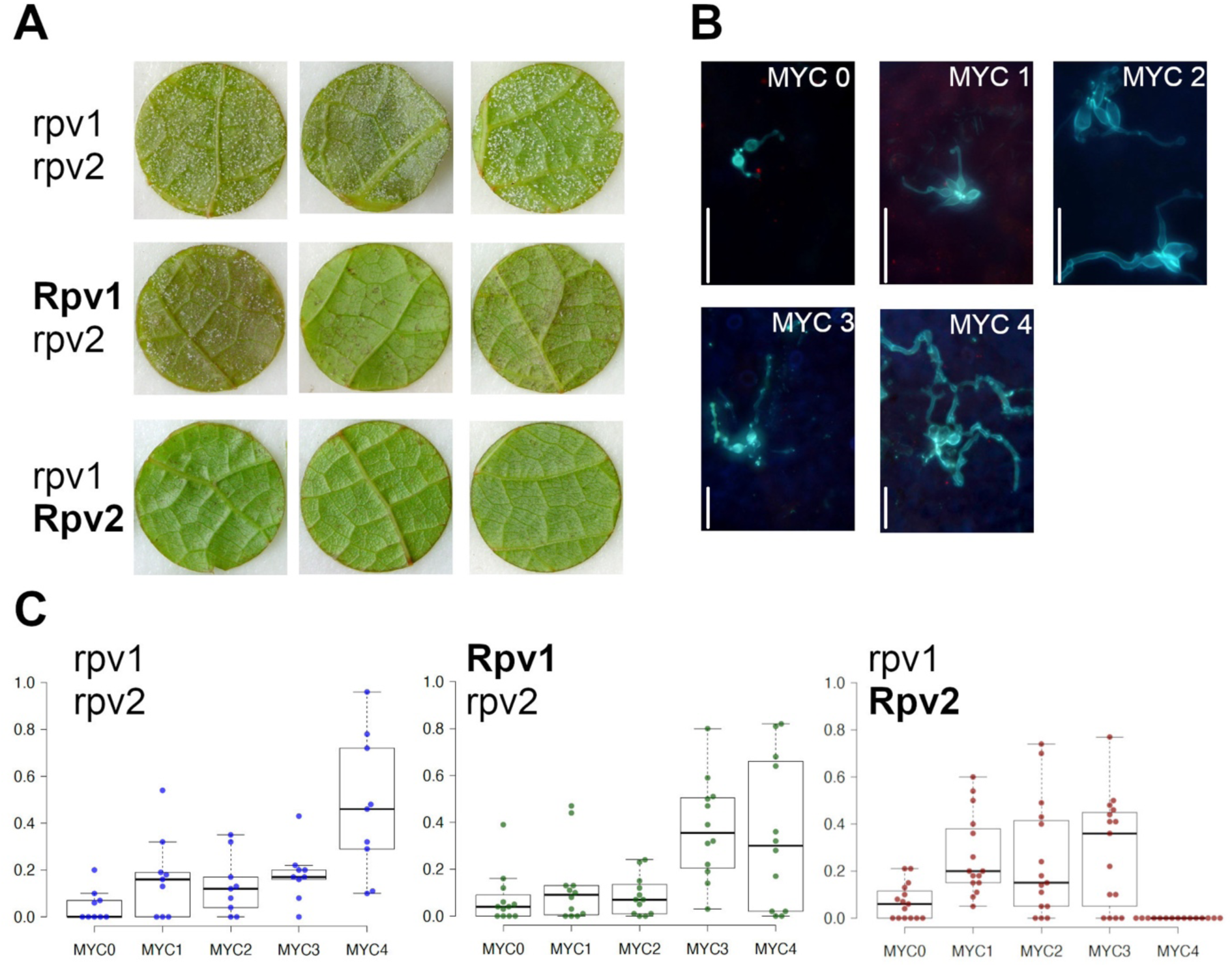
Effect of Rpv2 on *P. viticola* infection. (A) Macroscopic phenotype following *P. viticola* inoculation of genotypes from the pseudo- BC1 population carrying Rpv1 alone (middle), Rpv2 alone (bottom) or lacking both genes (top). Each disc corresponds to a genotype. (B) Referential of classes of intra-tissular pathogen development. Bars indicate 50 µm. (C) Percentage of observed infection events for each class of pathogen development in leaf discs from genotypes carrying Rpv1 alone (middle), Rpv2 alone (right) or lacking both genes (left). Each point corresponds to the average of five microscopic fields observed in one leaf disc. Three leaf discs were used per genotype. rpv1,rpv2: n = 9 (3 genotypes); **Rpv1**,rpv2: n = 12 (4 genotypes); rpv1,**Rpv2**: n = 15 (5 genotypes).

To gain insight into the effect of *Rpv2* on pathogen development, we performed microscopical observations at 48-hours post-inoculation, an early time of the infection when sporulation is not yet visible. We defined different classes of pathogen development based on the length and ramification of hyphae following aniline blue staining (Fig. 2B) and quantified them in plants carrying *Rpv1*, *Rpv2* or neither of them. Plants lacking both resistance loci presented mainly long, ramified hyphae; plants bearing *Rpv1* presented less developed hyphae; finally, in *Rpv2* plants infection events were blocked at earlier stages and never led to ramified hyphae, (Fig. 2C). Quantification the numbers of stomata, encysted spores and infected stomata did not reveal differences between each class of individuals, ruling out the possibility that any of these traits was responsible for the reduced *P. viticola* development in *Rpv2* plants (Fig. S4). In conclusion, our results indicate that *Rpv2* acts early upon infection, hindering pathogen development from the initial phases.

### Genetic mapping of Rpv2

To improve genetic mapping and move towards fine mapping of *Rpv2*, we used a pseudo-BC4 population, derived from a pseudo-BC1 individual carrying only *Rpv2*, consisting of 79 individuals (Delame et al. 2019).

Because *V. rotundifolia* has been reported to possess a gene for resistance to powdery mildew in the genomic region spanning *Rpv2*, plants were evaluated for resistance to downy and powdery mildews. Traits scored for both pathogens are presented in Table S2. The bimodal distribution of resistance classes for both pathogens (respectively OIV452 and OIV455 for downy and powdery mildew) in the population suggested monogenic resistance mechanisms (Fig. S5).

The population was then genotyped using SSR markers. Out of 328 markers tested, 148 were heterozygous in the resistant female parent (1171P), and 108 of them were assembled in 19 linkage groups. Four additional SSR markers were designed in the *Rpv2* genomic region on linkage group 18 based on the PN40024 12X reference genome sequence (Table S5). The resulting map covers 59% of the estimated 1166.8 cM reference integrated map distance and the average distance between markers is 6.89 cM (Fig. S6). Linkage group 18 was more densely covered, with an average distance between markers of 1 cM in the *Rpv2* region.

QTL analysis for downy mildew resistance revealed a strong QTL on linkage group 18 with all variables analysed (Table 1), which explained 93% of the phenotypic variance for OIV452. To assess if resistance to downy and powdery mildew were caused by the same gene or by different genes, the OIV452 and OIV455 variables were converted into discrete variables (OIV<6=susceptible; OIV≥6=resistant) and mapped as such. Resistance to powdery mildew (*Run2*) mapped at a telomeric position at 35 cM, while resistance to downy mildew (*Rpv2*) mapped at 27 cM (Fig. 3). These results show that *Rpv2* alone confers total resistance to downy mildew and that *Rpv2*-mediated resistance is independent from *Run2*.

**Figure 3.**
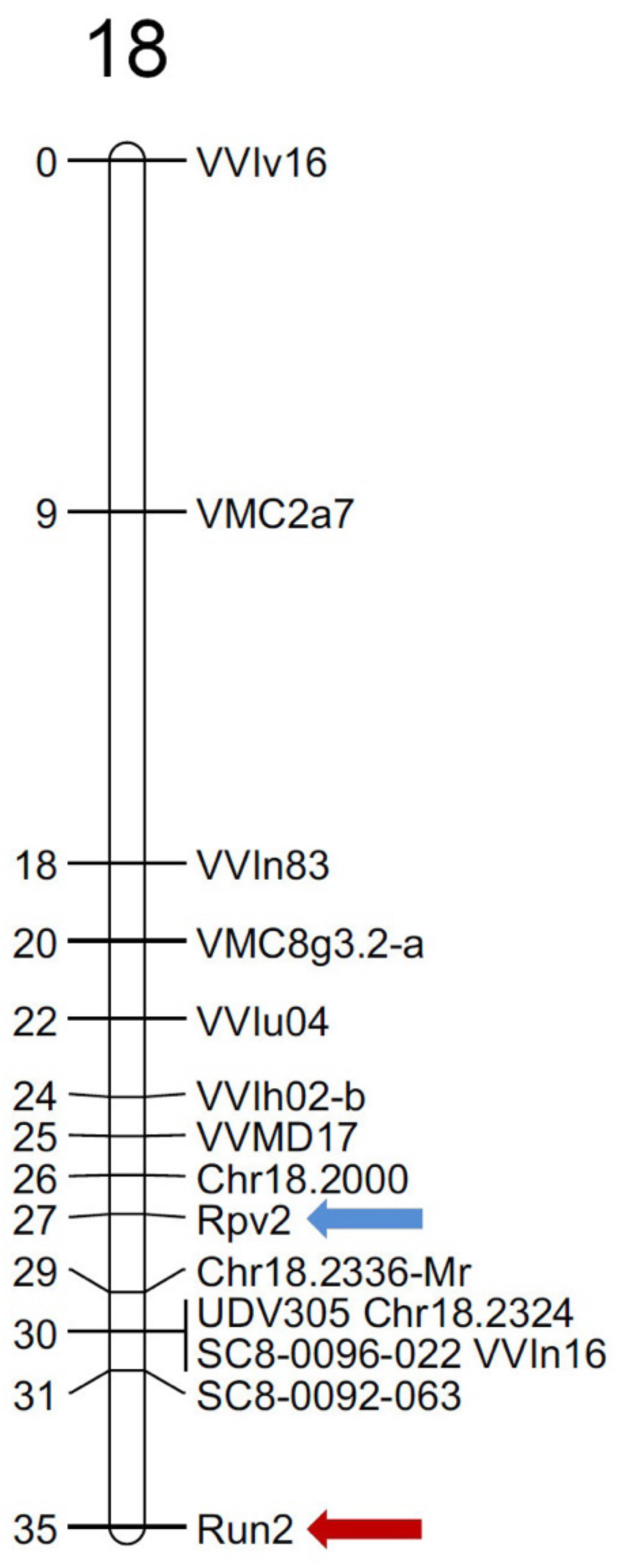
Mapping of *Rpv2* and *Run2.2* in the pseudo-BC4 population. Localisation of Rpv2 (blue arrow) and Run2.2 (red arrow) in linkage group 18 of the pseudo- BC4 genetic map. Semi-quantitative OIV452 and OIV455 variables for resistance respectively to downy and powdery mildew were converted into discrete variables and mapped as such. Distances are in cM.

**Table 1.** QTLs for resistance to *P. viticola* detected in the pseudo BC4 population.

| Trait | LG | Nearest marker | Peak (cM) | CI (cM) | LOD | % phenotypic variance |
| --- | --- | --- | --- | --- | --- | --- |
| OIV452 | 18 | Chr18.2000 | 26.79 | 25.79-28.79 | 44.45 | 93.0 |
| SA | 18 | Chr18.2000 | 26.79 | 24.55-29,50 | 19.55 | 70.6 |
| SNPB | 18 | Chr18.2000 | 26.79 | 21.92-30.79 | 9.28 | 45.3 |

**LG:** linkage group; **Peak:** position of the QTL peak; **CI:** QTL confidence interval.

### Fine mapping of *Rpv2*

*Rpv2* is located in an interval spanning roughly 3 Mb in the PN40024 12X reference genome. Our next aim was thus reducing the interval spanning *Rpv2*. The genotype for markers surrounding *Rpv2* of two individuals of the pseudo-BC4 population suggested that they were carrying recombination events inside the interval. Besides, in the process of *Rpv2* introgression, out of 100 individuals belonging to two pseudo-BC5 populations, four were identified as recombinant in the region. We sequenced using short-reads those six recombinant individuals and the parents of the pseudo-BC4 population (1771P and Nebbiolo). Two individuals non-recombinant in the *Rpv2* region (one resistant and one susceptible) were also sequenced to be used as controls. This information, together with available *V. rotundifolia* cv Trayshed sequence data, was used to perform *in silico* chromosome painting and identify the recombination points for each individual inside the *Rpv2* interval. Comparing the *V. rotundifolia*-derived chromosomal fragments with the resistance phenotype of each individual revealed that the interval spanning *Rpv2* was reduced to 780 kb (Fig. 4).

**Figure 4.**
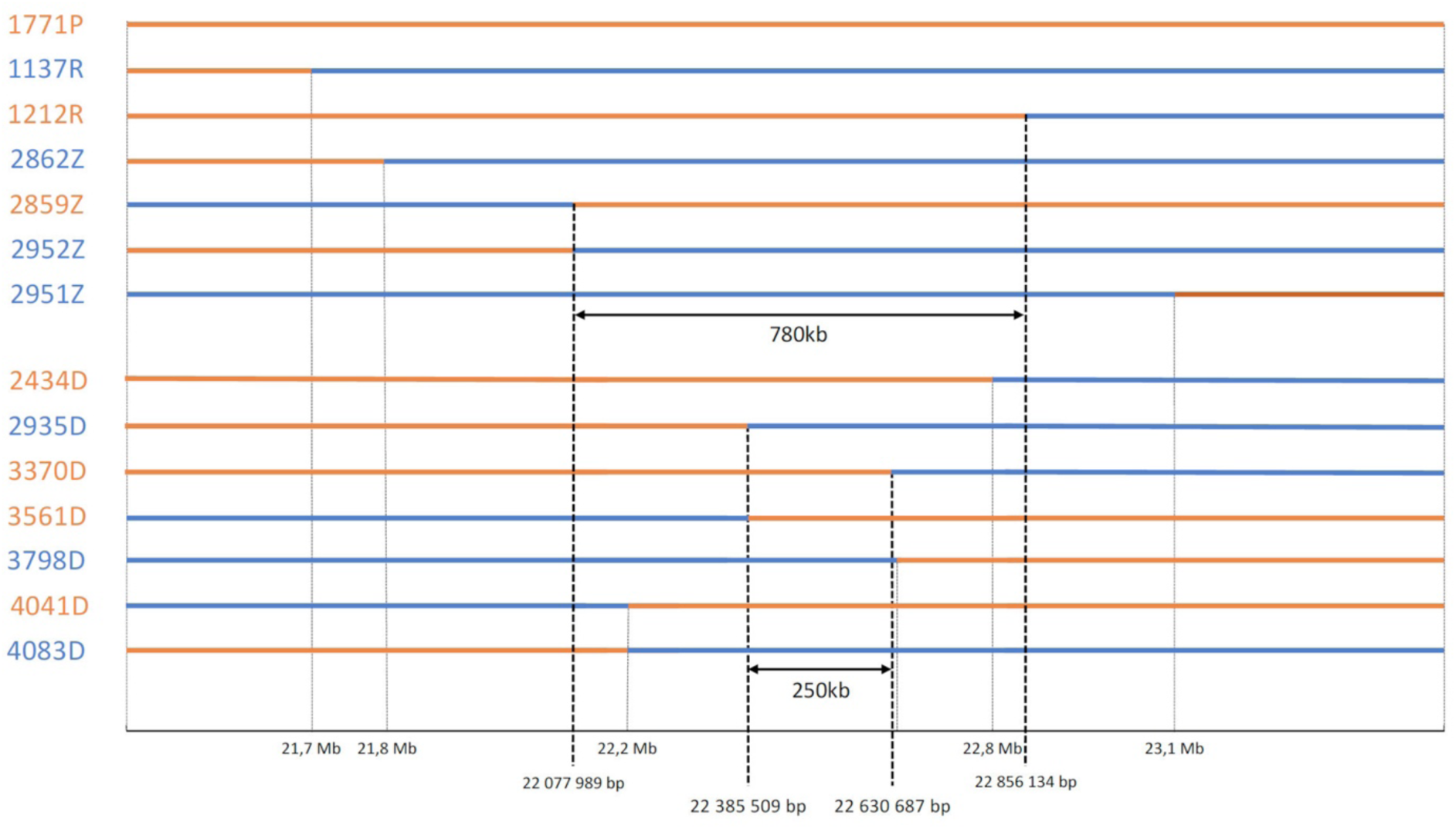
Fine mapping of *Rpv2*. *In silico* chromosome painting of recombinant individuals from different populations leading to the reduction of the *Rpv2*-containing genomic region to 780 kb and 250 kb. Names of genotypes in orange indicate resistant individuals and names in blue indicate susceptible individuals. Bars for each genotype represent the *Rpv2* genomic region from the chromosome inherited from the resistant donor. Orange indicates sequence from *V. rotundifolia* cv Trayshed and blue indicates sequence from *V. vinifera*. Positions correspond to the sequence of *V. rotundifolia* cv Trayshed Haplotype 1.

To further reduce the interval, we created two new populations obtained by self-pollination, one derived from the resistant parent of the pseudo-BC4 (1771P) and the second from one individual of the pseudo-BC4 population carrying *Rpv2*. To screen these new populations for recombinants inside the 780 kb interval, we designed molecular markers at both interval ends based on the comparison of the *V. rotundifolia* and *V. vinifera* sequences (Fig. S7). We screened 1508 individuals from both populations and identified 10 new recombinants, which were evaluated for downy mildew resistance. Short-read sequencing of seven recombinants allowed performing *in silico* chromosome painting, resulting in a further reduction of the *Rpv2* interval to 250 kb (Fig. 4).

### Genomic organization of the *Rpv2* region

To study the genomic organization of the *Rpv2* region, we took advantage of the availability of the diploid chromosome-scale assembly of the genome of *V. rotundifolia* cv Trayshed (Cochetel *et al*., 2021; Minio et al., 2022). Comparison of the *Rpv2* locus in the two haplotypes revealed that they are not identical (Fig. S8). To identify the haplotype carrying *Rpv2*, we used Illumina sequencing data from the BC4 individual 1126R, which is resistant and non-recombinant in the *Rpv2* locus. Reads were aligned on the two pseudomolecules of *V. rotundifolia* cv Trayshed and also compared to *V. vinifera* cv. Nebbiolo, which is the susceptible parent of the BC4 population. Nucleotide polymorphism analysis revealed that resistant haplotype from 1126R is identical to Haplotype 1 of *V. rotundifolia* cv Trayshed, and thus the *Rpv2* locus comes from Haplotype 1. Based on the automated annotation for the *V. rotundifolia* cv Trayshed genome (Cochetel et al. 2021) the 250 kb interval of Haplotype 1 encompassing *Rpv2* contains a cluster of NLR-type resistance genes. The high evolutionary dynamics of this kind of genes led us to perform our further ortholog analyses on the 780kb interval in order to have a clear overview of the locus.

It has been previously reported that *V. rotundifolia* cv Regale possesses a downy mildew resistance QTL localized on LG18 (Blanc et al. 2012), whose confidence interval encompasses the *Rpv2*-containing region in Trayshed. To understand the relationship between the locus in the two varieties, we generated a *V. rotundifolia* cv Regale genome assembly derived from PacBio Sequel sequencing (Table S6). Macro-synteny analysis using the Trayshed and Regale assemblies revealed the presence of sequences homologous to the Rpv2 locus in Regale contigs 69F, 730F and 165F, with contigs 69F and 165F showing homology with Trayshed haplotype 1 and contig 730F with Trayshed haplotype 2 (Fig. S9). These results suggest that the Rpv2 locus is conserved between the genomes of Regale and Trayshed.

Automatic annotation in the 780 kb interval from Trayshed Haplotype 1 predicted 19 genes belonging to different gene families (Cochetel et al. 2021). In order to cure the automatic annotation inside the *Rpv2* locus, we used RNA sequencing data that we generated from *V. rotundifolia* cv. Regale. Curation resulted in the identification of 25 genes in the interval, including a cluster of seven genes from the Nod-like receptor family (NLR) within the 250 kb interval (Fig. 5A, Table S7). The NLR cluster contains several genes encoding truncated proteins and only two genes (231970 and 232010) encode full-length proteins containing TIR, NB-ARC and LRR domains (Fig. 5B). Expression analysis of both genes in different conditions using the same transcriptomic data used for annotation revealed that they are both expressed, with gene 231970 displaying a distinctive level of expression in roots and in infected leaves compared to other conditions (Fig. 5C). These two full-length genes were selected as candidates for *Rpv2* resistance.

**Figure 5.**
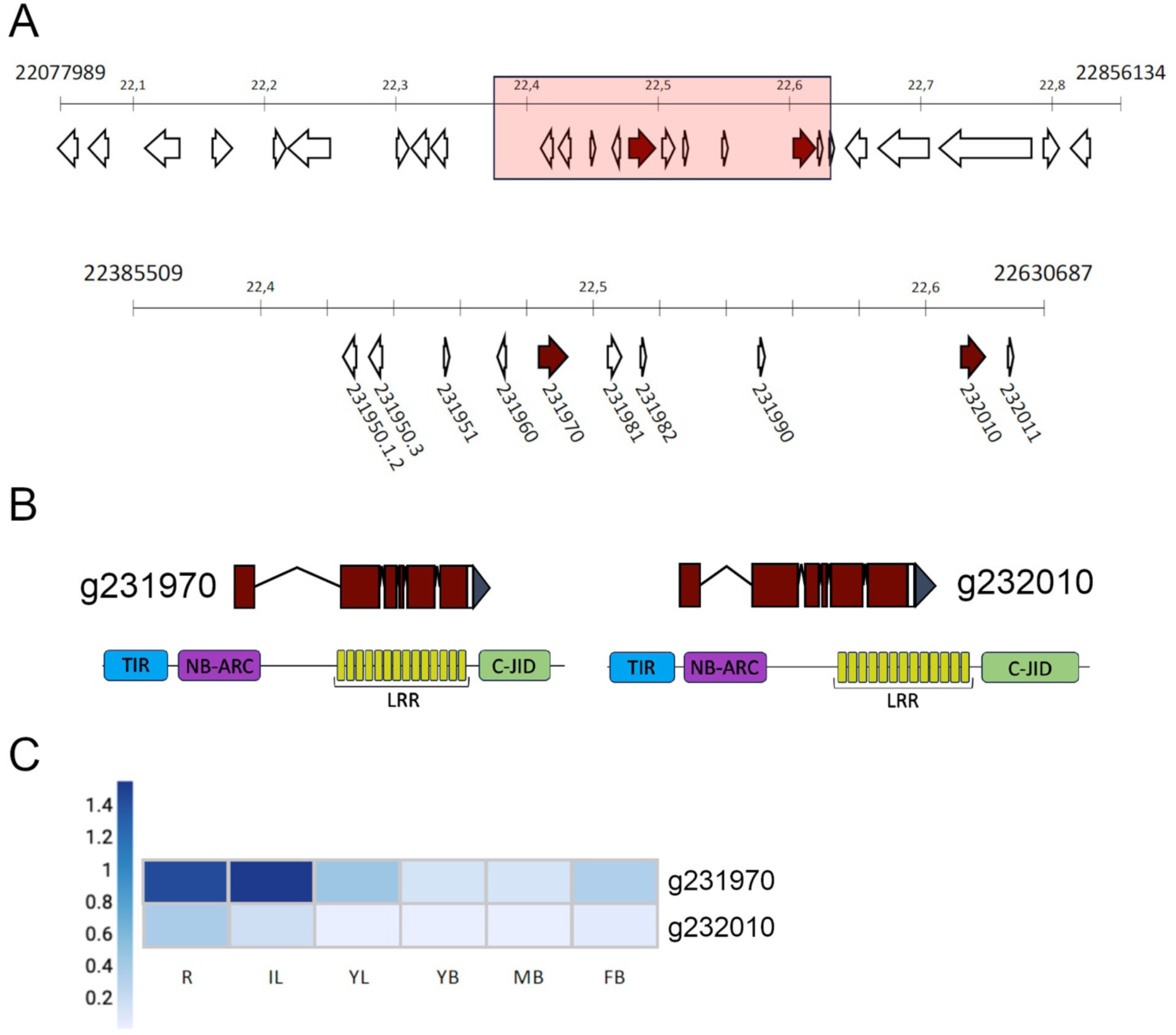
Genomic organization of the *Rpv2* locus in *V. rotundifolia* cv Trayshed. (A) Organization of the 780 kb genomic region containing *Rpv2*. Arrows represent genes. The 250 kb genomic interval containing Rpv2 is boxed and presented in detail below. The two complete NLRs found in the interval are colored in red. The genes present in the 250 kb interval are listed in Table S7. Physical locations are specified as the nucleotides on Trayshed haplotype 1 chromosome 18. (B) Intro-exon and protein domain organization of the two complete NLRs found in the 250 kb interval containing *Rpv2*. TIR: Toll/Interleukin-1 receptor; NB-ARC: nucleotide binding; LRR: leucine-rich repeat; C-JID: C-terminal jelly roll/Ig-like. (C): Heatmap of gene expression of both NLRs in different tissues of *V. rotundifolia* cv Regale. R: roots; IL: infected leaves; YL: young leaves; YB: young berries; MB: mature berries; FB: floral buds. Units presented as RPKM.

Next, we performed curated annotation of the 780 kb interval from Trayshed Haplotype 2 as well as of the Regale contigs belonging to the same interval. The analysis revealed eight genes coding for full-length TNLs, four from Trayshed Haplotype 2 and four from Regale. Comparison of the protein sequence of these eight TNLs with the two *Rpv2* candidates revealed that the TNLs present in contigs 69F and 165F from Regale are identical to the two full-length TNLs from Trayshed Haplotype 1, while all other genes show some level of divergence (Fig. S10). These results, along with the conservation at the micro-synteny level (Fig. 6A), reinforce the idea of the conservation of the *Rpv2* locus between Trayshed and Regale and suggest that *Rpv2* is also present in Regale.

**Figure 6.**
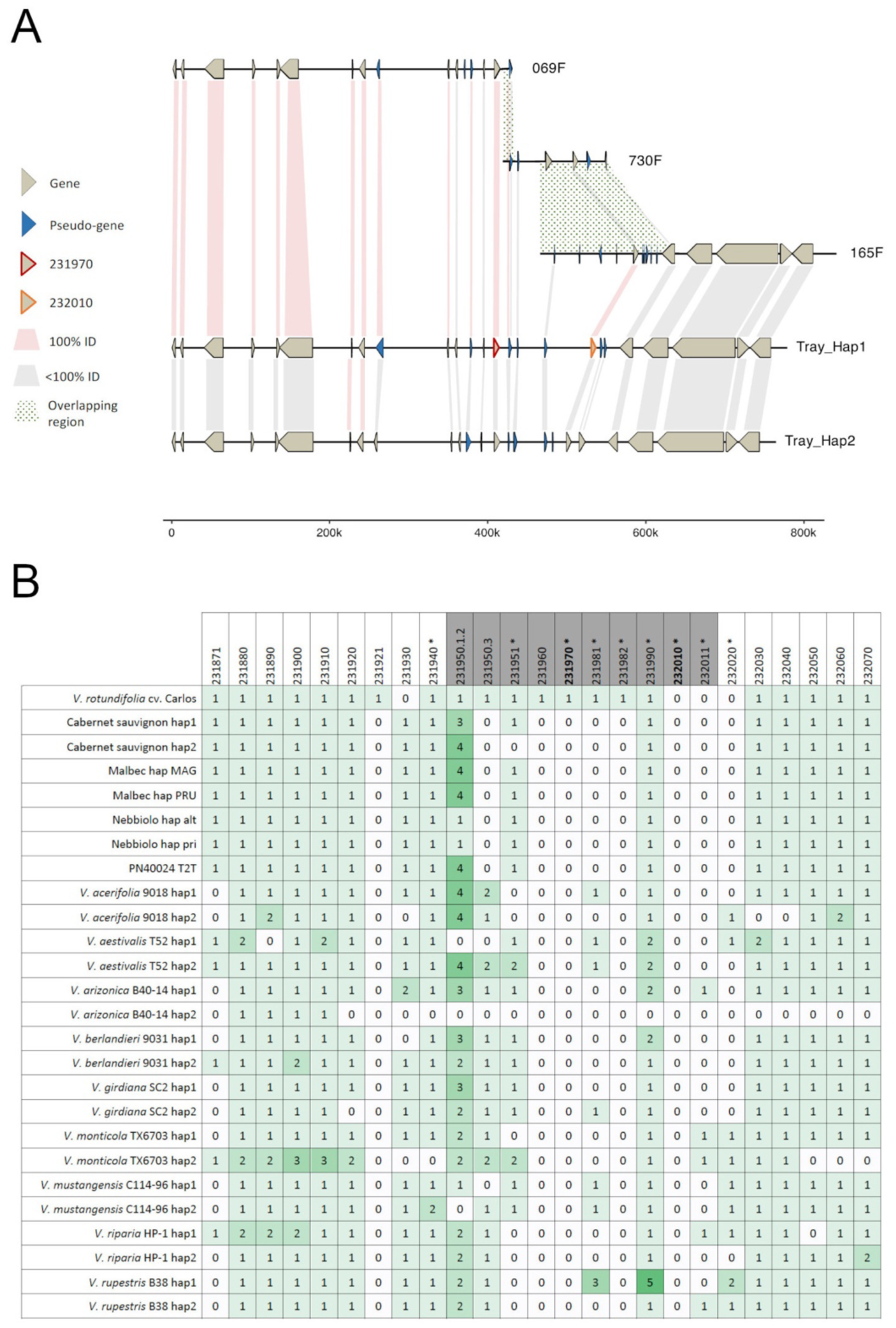
Synteny of the 780 kb *Rpv2* locus in various *Vitis* species. (A) Synteny analysis of curated gene annotations of *V. rotundifolia* cv. Trayshed haplotype 2 and contigs 069F, 730F and 165F of *V. rotundifolia* cv. Regale overlapping with the 780kb locus of *V. rotundifolia* cv. Trashed haplotype 1. Brown arrows correspond to complete genes and blue arrows to pseudogenes. Candidate genes are enhanced with red outline for 231970 and orange outline for 232010. Pink links correspond to the same protein sequence and grey links to an identity percentage between 90 and 100 %. Links with green dots represent overlapping regions with contigs of Regale. (B) Analysis of the Copy Number Variation of the 25 Trayshed Haplotype 1 genes in 14 *Vitis* genotypes (26 haplotypes). The number of copies is indicated with numbers from 0 to 5 with an intensity gradient of green colour. Genes highlighted in grey are present in the 250 kb interval. Genes with an asterisk belong to the TNL family. Genes in bold are the two candidate genes which are the only TNL with a full canonical structure.

To further confirm the *Rpv2* candidate genes, we performed comparative genomic analyses with diploid assembled genomes from *Vitis* species. Analysis of the micro-synteny between Trayshed Haplotype 1 and 14 *Vitis* genotypes at the 780 kb interval revealed sequence conservation at both extremities of the interval and divergence at the internal region of 250 kb containing Rpv2 (Fig. 6B). Orthologues of the genes localized at the left extremity of the 250 kb interval (231950.1.2, 231950.3 and 231951, which is a pseudogene with NB and LRR domains) were found in the genomes of several *Vitis* species, as well as 231990 (pseudogene with LRR domain). Orthologues of the other genes of the 250 kb interval were either absent or present in only few genomes. Thus, we did not find orthologues of the two *Rpv2* candidates (231970 and 232010) in any of the assembled genomes. The only exception is an ortholog of 231970 that was found for *V. rotundifolia* cv. Carlos but the identity percentage at the protein level was only 90.7%. These results suggest that both genes are not found outside *V. rotundifolia* and reinforce their condition as *Rpv2* candidate genes.

## DISCUSSION

Here we described the identification and characterization of *Rpv2*, a gene for extreme resistance to grapevine downy mildew derived from *V. rotundifolia* cv Trayshed. Microscopical analysis showed that *Rpv2*-mediated resistance results in pathogen arrest at the early stages of the infection cycle. Genetic and physical mapping led to the restriction of *Rpv2* into a 250 kb genomic region that notably contains two resistance genes of the NLR type. Comparative genomics revealed that orthologs of both NLRs could not be found in the other *Vitis* species that were analysed.

*Rpv2* differs from other *Rpv* genes in that the resistance it confers is total. Indeed, while for all reported *Rpv* genes the resistance is associated to macroscopic cell death and some level of sporulation, *Rpv2*-mediated resistance does not allow sporulation and it is often associated to microscopic cell death that is limited to stomata companion cells and a few surrounding cells. Microscopic analysis revealed that the *Rpv2* resistance phenotype is not caused by an effect on the ability of the *P. viticola* spores to encyst and enter into the grapevine leaves, since the number of encysted spores and infected stomata per surface unit was the same in plants carrying *Rpv2* than in plants lacking it. Conversely, the microscopic analysis showed that Rpv2 stopped *P. viticola* infection at the early stages of pathogen development inside plant leaves, following pathogen growth. The phenotype of the *Rpv2*-mediated resistance thus suggests that activation of the resistance requires a contact between *P. viticola* and the leaf mesophyll cells, most likely following the formation of the first haustoria, and hence that resistance is mediated by a receptor-like protein.

The *Rpv2* locus was reduced to a genomic region of chromosome 18 of 250 kb in Haplotype 1 of the genome assembly of *V. rotundifolia* cv Trayshed. Curated annotation of the region showed that it contains a cluster of NLR-like genes, including two complete genes (with TIR, NBS and LRR domains), named 231970 and 232010. Genes orthologous to 231970 and 232010 from Haplotype 2 of Trayshed were slightly different at the protein level (96.52% and 91.27% identity, respectively). Interestingly, the syntenic region of the genome assembly of *V. rotundifolia* cv Regale contained two TNLs that were 100% identical at the protein level to 231970 and 232010. Regale has been reported to carry a locus for resistance to downy mildew on linkage group 18, in the same region as *Rpv2* (Blanc et al. 2012). *V. rotundifolia* grows in an area of high pathogen pressure, considered to be the *P. viticola* centre of origin; it would be thus expected that functional resistance genes against *P. viticola* are subjected to selection pressure and conserved among natural populations. In this context, the fact that genes 231970 and 232010 are 100% identical between Trayshed and Regale makes them strong candidates for being responsible for the resistance found in both varieties, and thus strong candidates for *Rpv2*.

Results of comparative genomics supported the role of 231970 and 232010 as *Rpv2* candidates. While we could find the orthologues for some other genes of the region, we didn’t find orthologues of 231970 and 232010 in the assembled genomes of any of the *Euvitis* species analyzed, indicating that they may be *V. rotundifolia* – specific. An ortholog of 231970 was found in the assembled genome of *V. rotundifolia* cv. Carlos with a low identity percentage suggesting that the candidate alleles were not found in Carlos. However, this does not call into question our hypothesis of 231970 and 232010 being *Rpv2* candidates. On one hand, to our knowledge, despite Carlos resistance to downy mildew, it has never been demonstrated that Carlos carries *Rpv2*. On the other hand, the genome assembly of Carlos (Huff et al. 2023) does not resolve the two haplotypes and 231970 and 232010 genes may well be present on the non- assembled one. Further support for the role of 231970 and 232010 as *Rpv2* candidates came from transcriptome analysis, which confirmed that both candidates are expressed, indicating that both must be functional. Further research will be required to determine which of the genes, if not both, is responsible of the *Rpv2*-mediated resistance.

*V. rotundifolia* cv Trayshed carries a gene for resistance to powdery mildew, named *Run2.2*, which is also located in the distal part of chromosome 18 (Riaz et al. 2011). Mapping the resistance to downy and powdery mildew in the pseudo-BC4 population allowed us to genetically separate *Rpv2* and *Run2.2*. Comforting our results, we located *Run2.2* at the same position as previously reported (Riaz et al. 2011). Putting together our results with previous reports (Massonnet et al. 2022), *Rpv2* and *Run2.2* are both contained in the same haplotype of the *V. rotundifolia* cv Trayshed genome, separated by at least 10 Mb of sequence. This observation has implications for breeding: on one hand, the fact that both genes are in the same haplotype makes possible to transfer them together in the introgression process; however, on the other hand, introgressing together the resistance to both pathogens from *V. rotundifolia* Trayshed would imply dragging a big fragment of *V. rotundifolia* genome (Delame et al. 2019). Understanding the phenotype associated to this linkage drag will be helpful to improve the efficiency of the introgression of both genes.

In conclusion, here we showed that *Rpv2* belongs to the NBS-LRR class of disease resistance proteins. The strong effect of *Rpv2* on pathogen development suggests that it detects a *P. viticola* effector expressed early upon infection and that its activation leads to a rapid and efficient resistance response. The total resistance conferred by *Rpv2* makes it interesting for breeding, because the absence of pathogen sporulation will result in an important reduction of pathogen pressure; however, due to its NLR nature it may be easily overcome by the pathogen, and thus it should be used in combination with other resistance genes to avoid resistance-breaking. The identification of the corresponding *Avr* gene and the study of its variability in *P. viticola* populations will shed light on the risk of *Rpv2* resistance breaking and also provide a tool for monitoring the eventual arising of virulence in pathogen populations. Finally, it is worth noting that since there have been no reports of *P. viticola* isolates overcoming the resistance from *V. rotundifolia*; it remains possible that combining *Rpv1* with *Rpv2* provides durable resistance to downy mildew.

## ACKNOWLEDGEMENTS

We thank Gisele Butterlin for technical assistance in RNA extraction, Florent Waltz for assistance in phenotyping and Anne Alais for technical assistance in genotyping. We are indebted to the UEAV at INRAE Colmar for their assistance in the production and maintenance of plant material. The authors thank Aurelie Berard and Isabelle Le Clainche (EPGV Unit, INRAE Genomic Platform) for performing sequencing. We acknowledge the support of the Fondation Jean Poupelain through the HealthyGrape2 project.

## AUTHOR CONTRIBUTIONS

DM, CR and PM planned and designed the research. LM, EP, SW, LS, MD and SC performed experiments. LM, SB, BA, GB and CR performed bioinformatic analyses. AV and VD collected data. EP, SW, DM and PM analyzed experimental data. DM, CR and PM supervised research. LM, EP, DM, CR and PM wrote the manuscript. All authors read and approved the final manuscript.

## DATA AVAILABILITY

All other data is available as supplementary material.

