## Supplemental Figures S1 to S10 and Supplemental Tables S1 to S7 for "*Rpv2* is part of a cluster of NLRs specific to *Vitis rotundifolia* and confers extreme resistance to grapevine downy mildew"

**Supplemental Figure S1.** Pedigree of the populations used in this study.
Green shading indicates *V. vinifera* varieties (only the name of the variety is indicated for clarity). Orange shading indicates populations. Populations used in this study are outlined. Individuals used as parents of the populations relevant for this study are indicated in red.

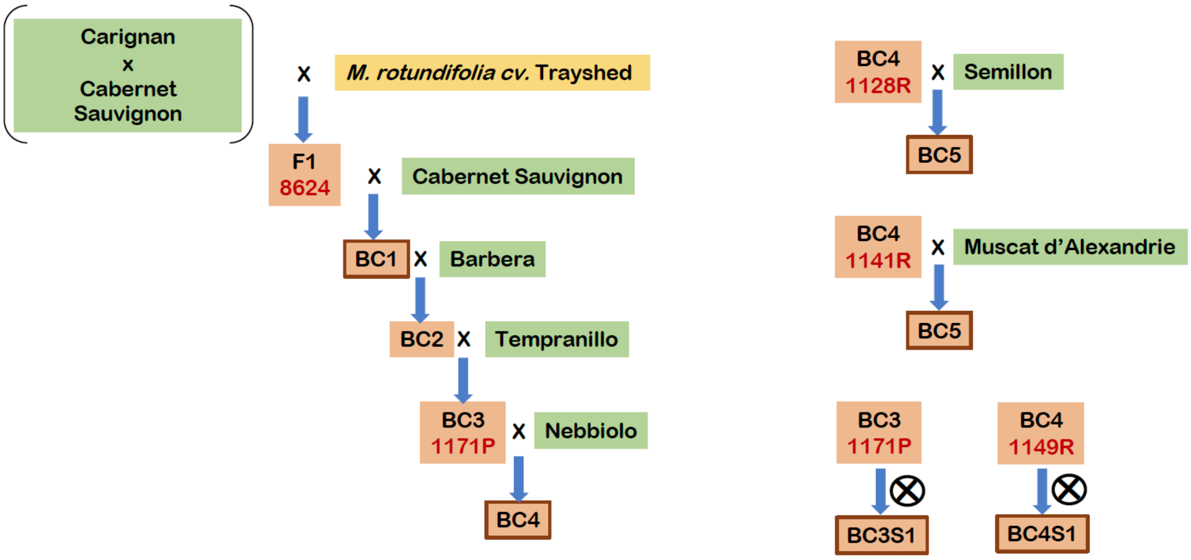

**Supplemental Figure S2.** Distribution of traits for downy mildew resistance in the pseudo-BC1 population. OIV452: symptom-based semi-quantitative score, N: presence of cell death; S: sporulation; SQ: sporulation intensity; NBSCP: sporangia per mL; DS: percentage of sporulating discs; NSTO: presence of stomatic necrosis. Ordinates represent number of individuals. Results for each individual correspond to the average of two biological repeats. Details about variables and scoring of are presented in Supplementary Table 3.

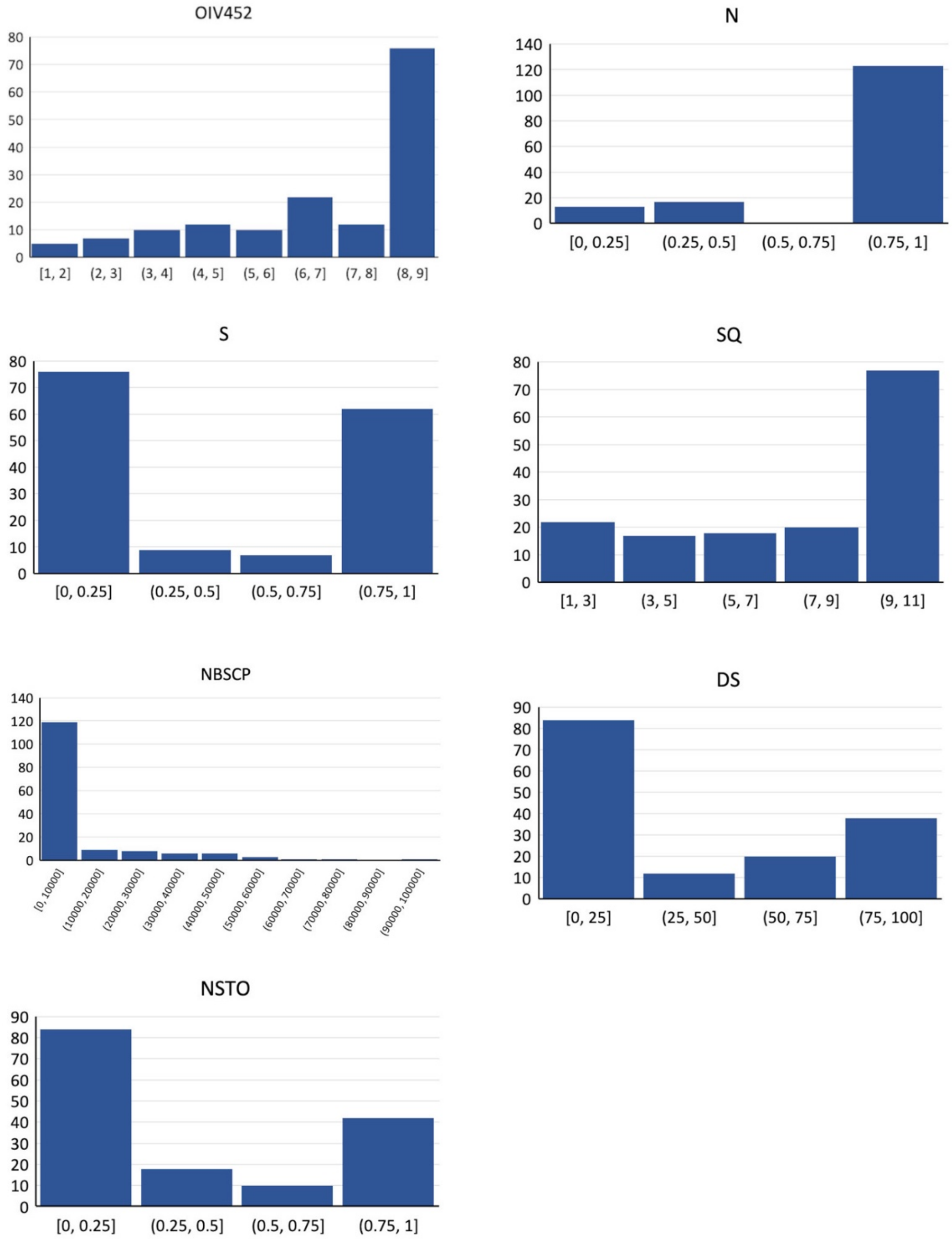

**Supplemental Figure S3.** Parental and consensus genetic maps of the pseudo-BC1 population. 8624: female resistant parent. CS: Cabernet sauvignon, male parent.

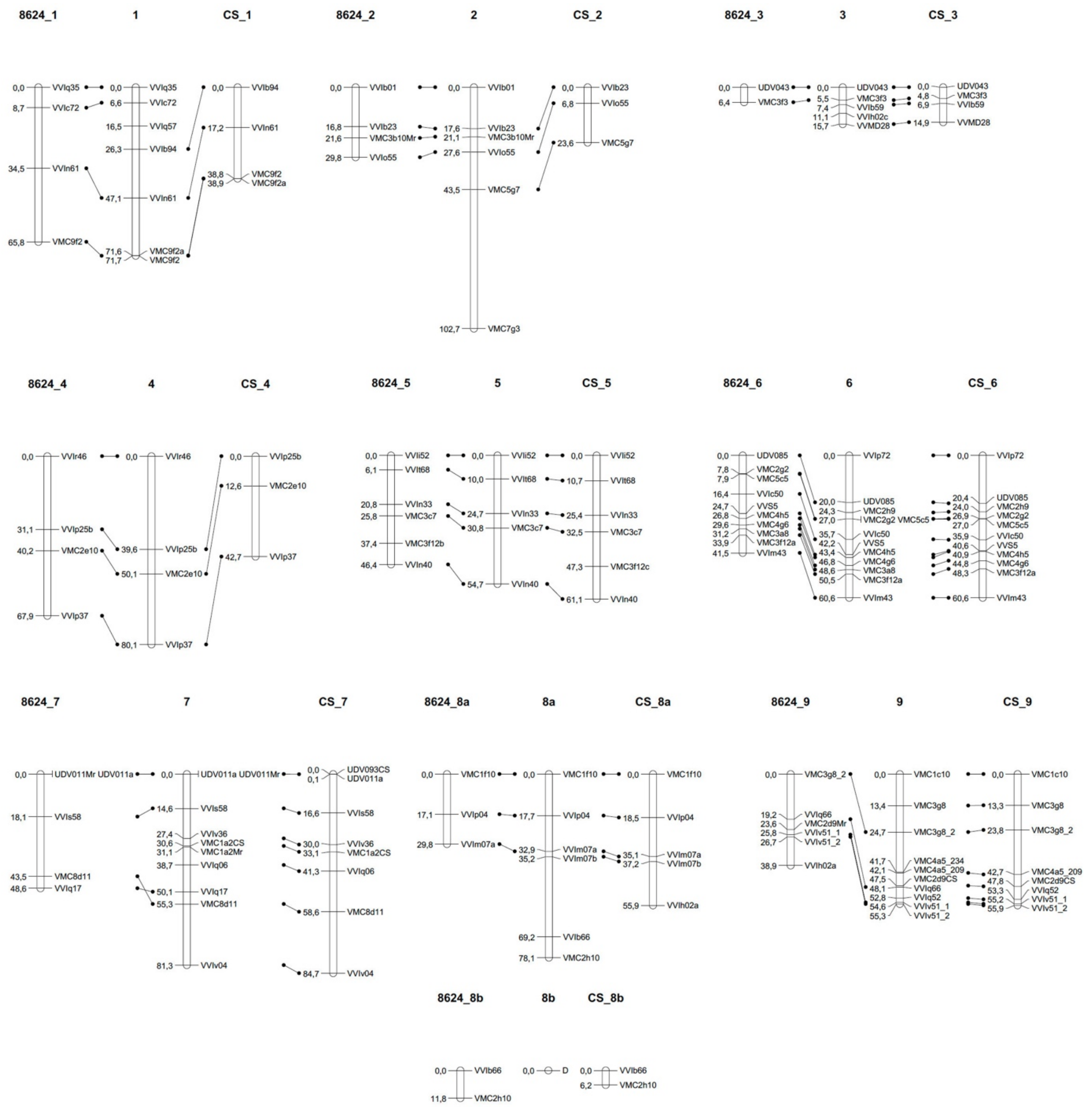

**Supplemental Figure S3** (cont)

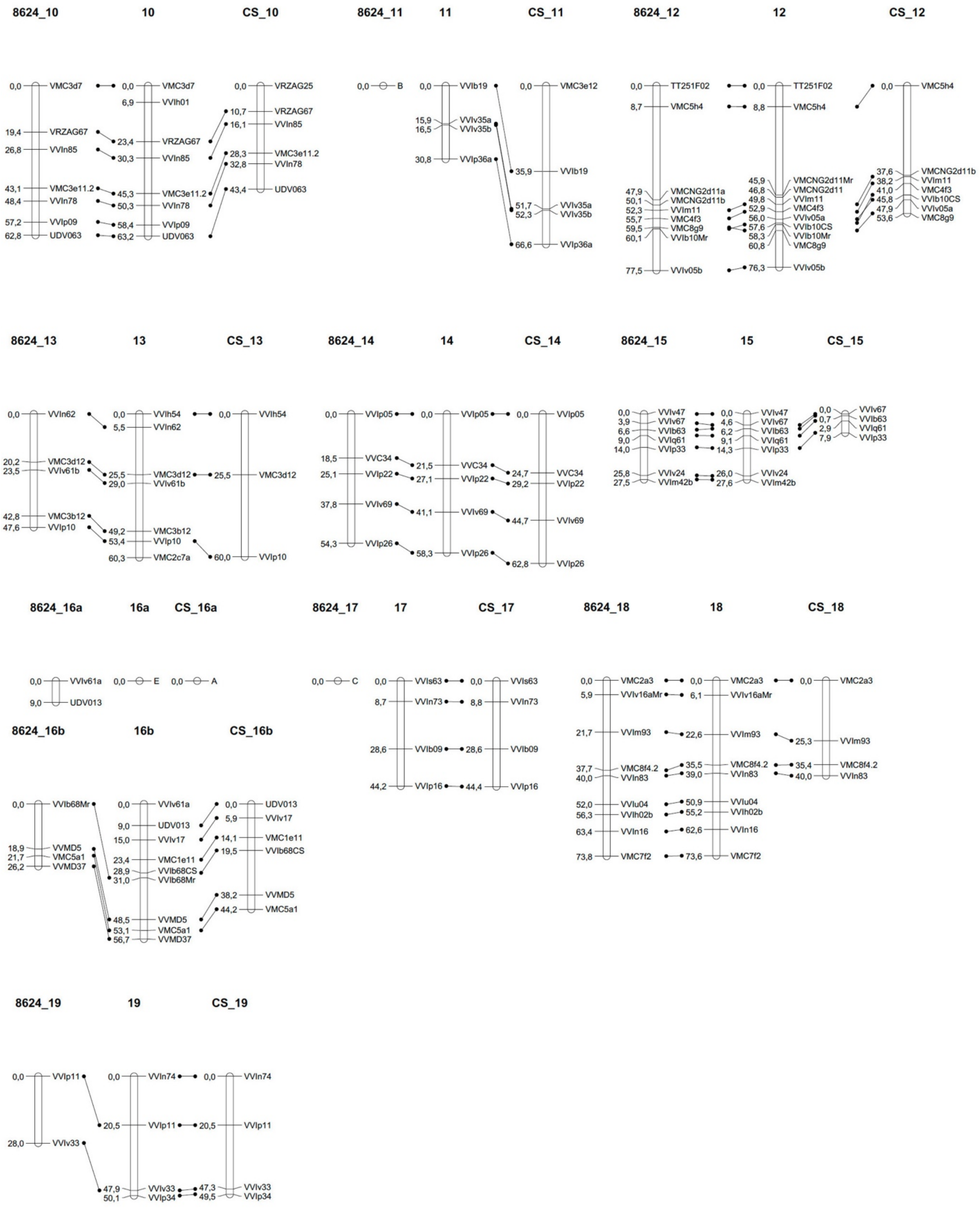

**Supplemental Figure S4.** Quantification the numbers of stomata, encysted spores and infected stomata on inoculated leaf discs.

Number of stomata, encysted spores and infected stomata present in leaf discs from genotypes carrying *Rpv1* alone (middle), *Rpv2* alone (right) or lacking both genes (left). Each point corresponds to the average of five microscopic fields observed in one leaf disc. Three leaf discs were used per genotype. Rpv1,rpv2: n = 9 (3 genotypes); **Rpv1**,rpv2: n = 12 (4 genotypes); rpv1,**Rpv2**: n = 15 (5 genotypes).

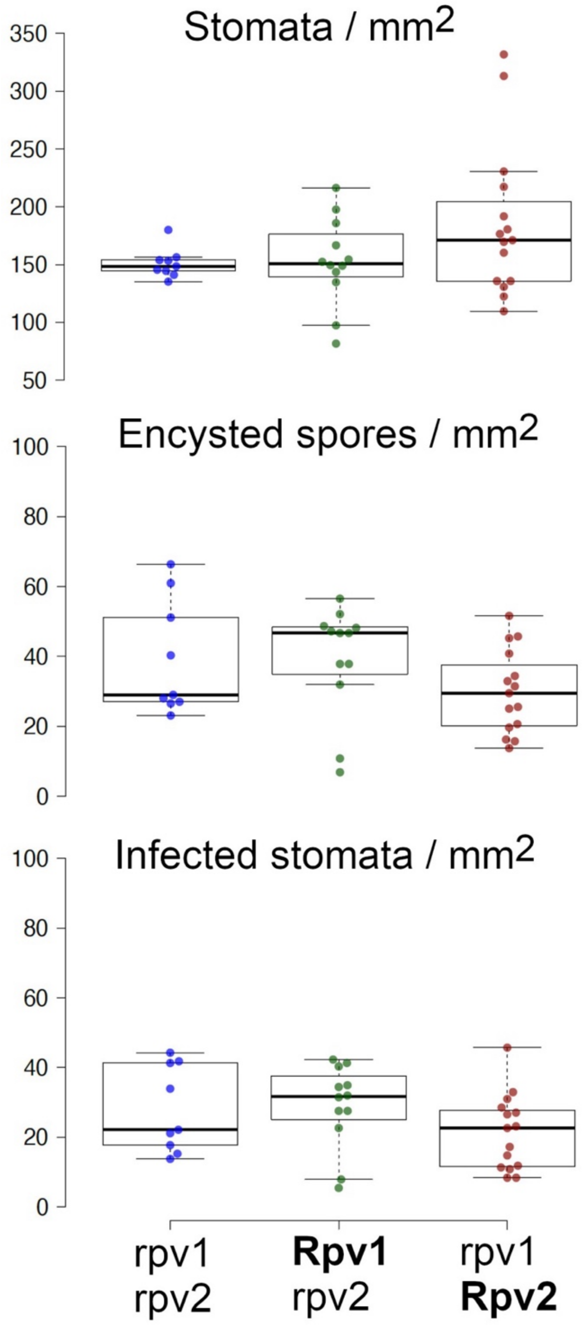

**Supplemental Figure S5**. Distribution of traits for downy and powdery mildew resistance in the pseudo-BC4 population.
5A) Downy mildew, (B) powdery mildew. OIV452: symptom-based semi-quantitative score for downy mildew resistance; SA: sporulating area; SPNB: sporangia per ml; OIV455: symptom-based semi-quantitative score for powdery mildew resistance. For downy mildew, results for each individual correspond to the average of two biological repeats for OIV452 and SA, and one repeat for SPNB. For powdery mildew, results for each individual correspond to one repeat for OIV455. Details about variables and scoring of are presented in Supplementary Table 5.

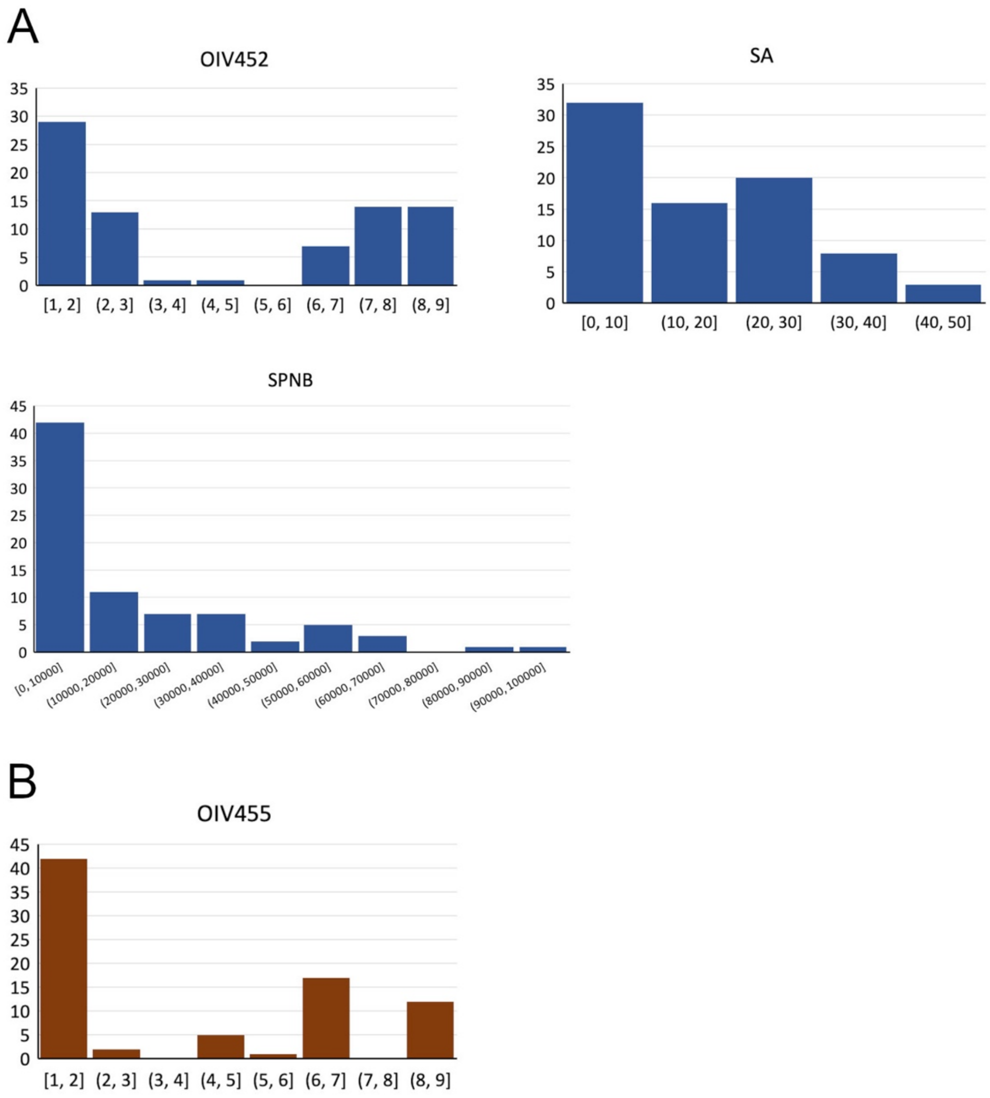

**Supplemental Figure S6.** Genetic map of the resistant parent of the pseudo-BC4 population.

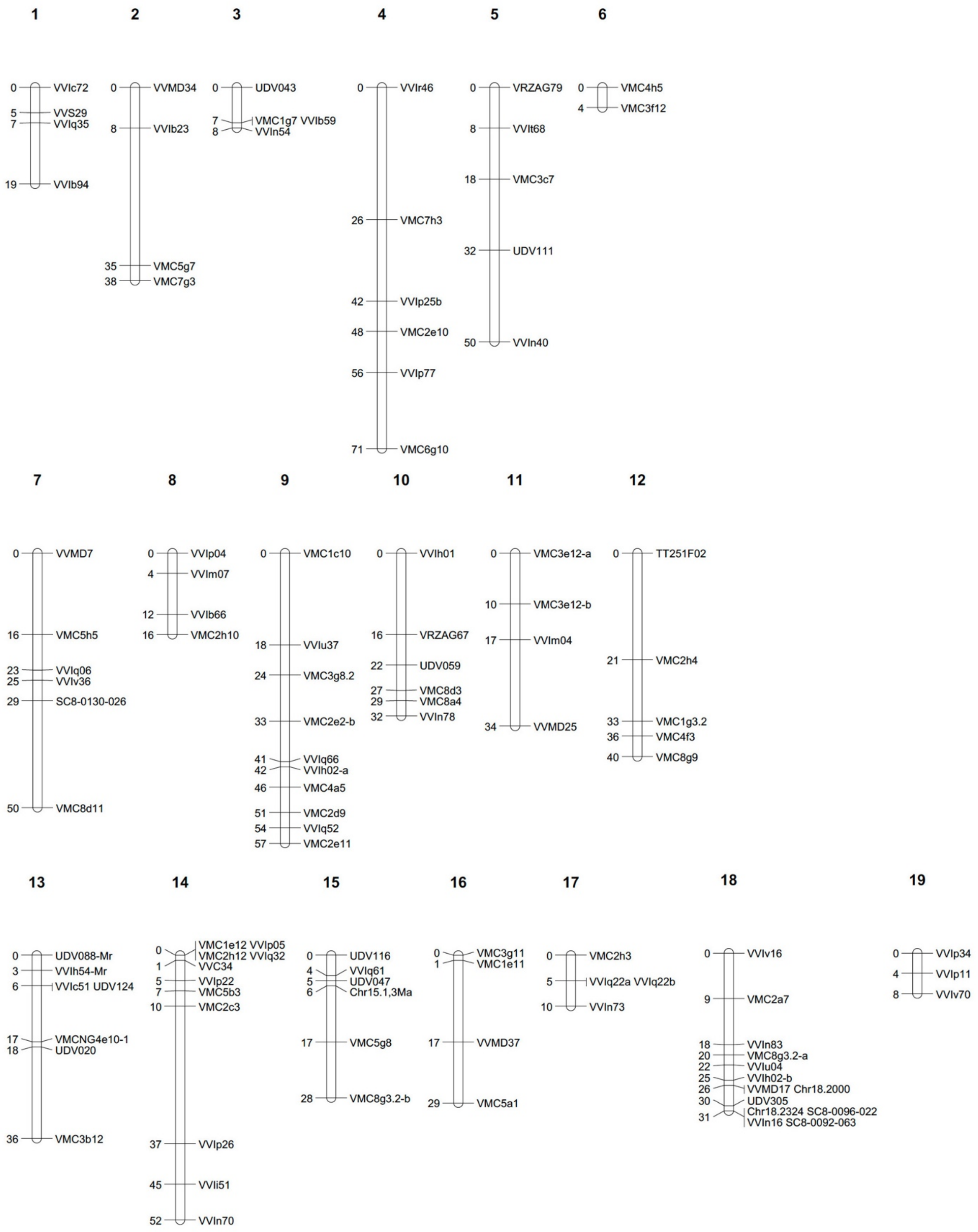

**Supplemental Figure S7.** Design of molecular markers inside the 780 kb interval.
Markers were designed around positions 22.0 MB (M220C, top) and 22.8 Mb (M228C, bottom). Images show the alignment of Illumina reads of V. rotundifolia cv Trayshed (bottom), Nebbiolo (susceptible parent of the pseudo-BC4 population, middle) and genotype 1771P (resistant parent of the pseudo-BC4 population, top) on Haplotype 1 of the genome assembly of *V. rotundifolia* cv Trayshed. Primers were designed to encompass the indels. Red arrows indicate the position of the primers. Primer sequences are shown. Note different scales between the two images.

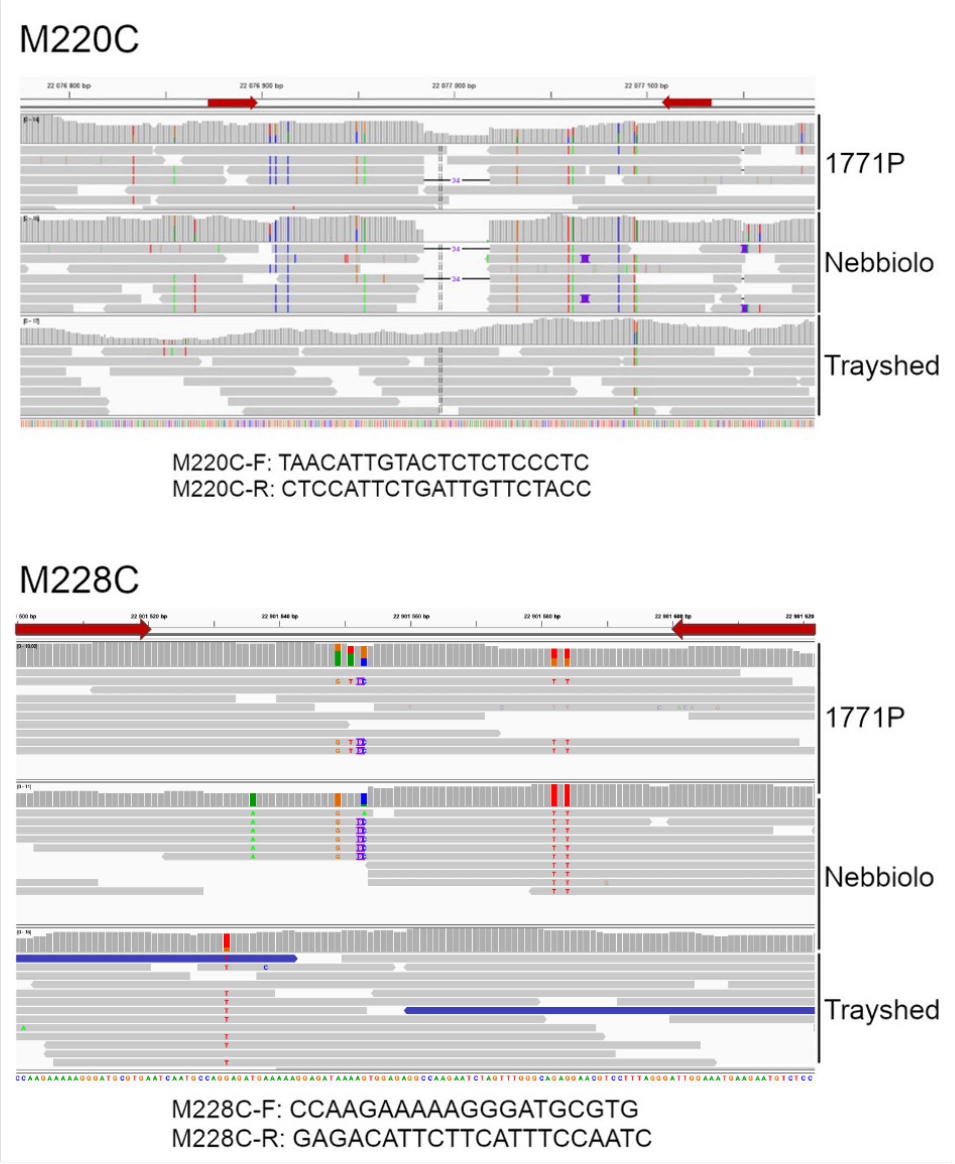

**Supplemental Figure S8.** Alignment of the *Rpv2* genomic region from the two haplotypes of the *V. rotundifolia* cv Trayshed genome.

Dot Plot showing the alignment of the 780 kb region encompassing *Rpv2* from the two haplotypes of the *V. rotundifolia* cv Trayshed genome. Blue represents forward alignments and purple reverse ones.

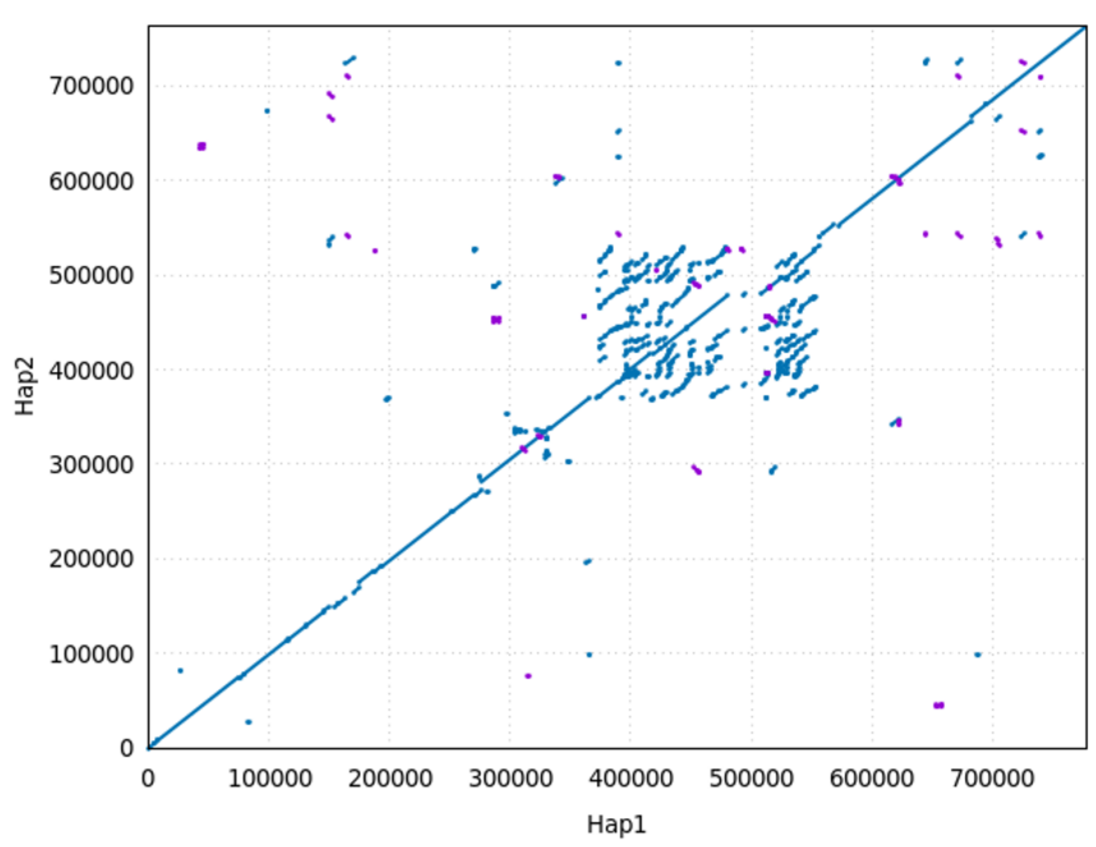

**Supplemental Figure S9.** Alignment of the genome assembly of *V. rotundifolia* cv Regale to the *Rpv2* genomic region from the two haplotypes of the *V. rotundifolia* cv Trayshed genome.
Dot Plot showing the alignment of the genome assembly of *V. rotundifolia* cv Regale to the 780 kb region encompassing *Rpv2* from the two haplotypes of the *V. rotundifolia* cv Trayshed genome. Stretches of 50000 X were added between contigs of Regale to distinguish them clearly.

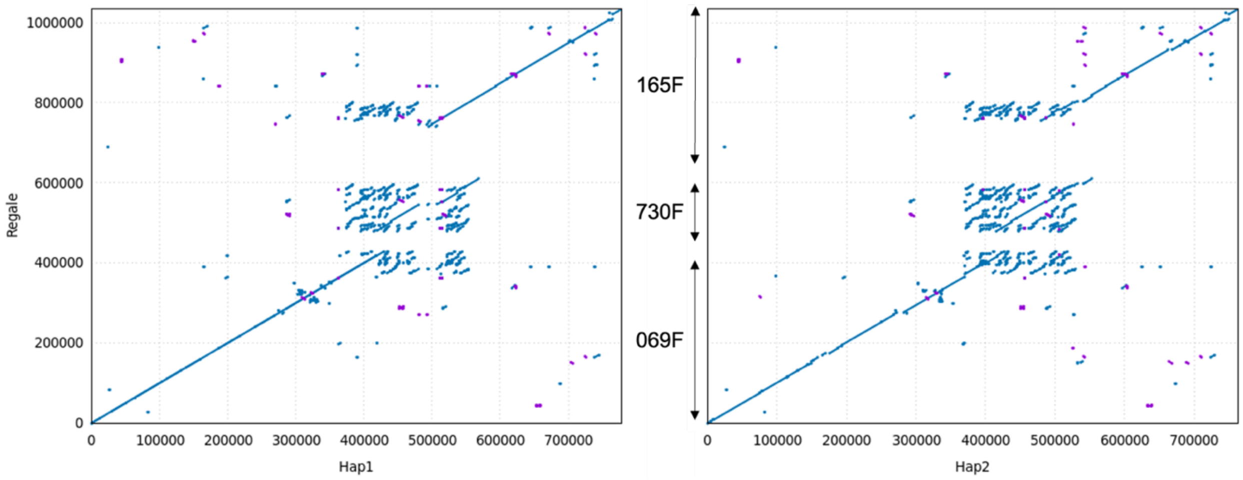


**Supplemental Figure S10**. Similarity matrix of complete NLR proteins found in the Rpv2 genomic region.
Similarity matrix of complete NLR proteins found in the 780 kb genomic region encompassing *Rpv2* from the genome assemblies of *V. rotundifolia* cv Trayshed (Haplotypes 1 and 2) and Regale. Note the 100% protein identity between the two complete NLRs from the 250 kb interval from Trayshed Haplotype 1 (232010 and 231970) with their orthologues from Regale.

| | g232010_Tray_hap2 | **g232010_Tray_hap1** | g232010a_Reg_ctg165F | g232010b_Reg_ctg730F | **g231970_Tray_hap1** | g231970_Reg_ctg069F | g231970_Tray_hap2 | g472510b_Tray_hap2 | g472510a_Tray_hap2 | g472510_Reg_ctg730F |
|---|---|---|---|---|---|---|---|---|---|---|
| g232010_Tray_hap2 | 100 | 91,27 | 91,27 | 91,36 | 90,5 | 90,5 | 89,49 | 77,38 | 74,1 | 75,75 |
| **g232010_Tray_hap1** | 91,27 | 100 | 100 | 99,45 | 92,74 | 92,74 | 92,12 | 77,66 | 71,73 | 74,74 |
| g232010a_Reg_ctg165F | 91,27 | 100 | 100 | 99,45 | 92,74 | 92,74 | 92,12 | 77,66 | 71,73 | 74,74 |
| g232010b_Reg_ctg730F | 91,36 | 99,45 | 99,45 | 100 | 92,9 | 92,9 | 92,29 | 77,76 | 72,65 | 74,83 |
| **g231970_Tray_hap1** | 90,5 | 92,74 | 92,74 | 92,9 | 100 | 100 | 96,52 | 78,36 | 70,97 | 74,91 |
| g231970_Reg_ctg069F | 90,5 | 92,74 | 92,74 | 92,9 | 100 | 100 | 96,52 | 78,36 | 70,97 | 74,91 |
| g231970_Tray_hap2 | 89,49 | 92,12 | 92,12 | 92,29 | 96,52 | 96,52 | 100 | 78,62 | 70,97 | 74,83 |
| g472510b_Tray_hap2 | 77,38 | 77,66 | 77,66 | 77,76 | 78,36 | 78,36 | 78,62 | 100 | 82,44 | 82,04 |
| g472510a_Tray_hap2 | 74,1 | 71,73 | 71,73 | 72,65 | 70,97 | 70,97 | 70,97 | 82,44 | 100 | 90,05 |
| g472510_Reg_ctg730F | 75,75 | 74,74 | 74,74 | 74,83 | 74,91 | 74,91 | 74,83 | 82,04 | 90,05 | 100 |

**Supplemental Table S1**. Variables used to assess the resistance level to downy mildew in the pseudo-BC1 population.

| Variable | Description | Scoring |
|---|---|---|
| OIV452 | Symptom-based semi-quantitative score of downy mildew resistance according to criteria of the Office International de la Vigne et du Vin (OIV; Anonymous 1983) | From 1 (very susceptible) to 9 (totally resistant) |
| S | Visual scoring of sporulation | 0 (absence) or 1 (presence) |
| SQ | Visual semi-quantitative scoring of sporulation intensity | From 1 (very dense) to 9 (not dense). Absent: 11. |
| DS | Percentage of sporulating leaf discs | From 0 to 100% |
| NBSCP | Number of sporangia per mL produced by leaf discs measured with a Z2 Coulter cell counter (Beckman Coulter). | Quantitative |
| N | Visual scoring of cell death | 0 (absence) or 1 (presence) |
| NSTO | Scoring of stomatal and peri-stomatal necrosis with a stereomicroscope (0.04-0.1 mm black necrotic points) | 0 (absence) or 1 (presence) |

**Supplemental Table S2**. Variables used to assess the resistance level to downy and powdery mildews in the pseudo-BC4 population.

| Pathogen | Variable | Description | Scoring |
|---|---|---|---|
| Downy mildew | OIV452 | Symptom-based semi-quantitative score of downy mildew resistance according to criteria of the Office International de la Vigne et du Vin (OIV; Anonymous 1983) | From 1 (very susceptible) to 9 (totally resistant) |
| Downy mildew | SA | Sporulating area assessed by image analysis | From 0 –to 100% |
| Downy mildew | SPNB | Number of sporangia per mL measured with a cell counter, Scepter™ 2.0 Cell Counter (Merck Millipore) | Quantitative |
| Powdery mildew | OIV455 | Symptom-based semi-quantitative score of powdery mildew resistance according to criteria of the Office International de la Vigne et du Vin (OIV; Anonymous 1983) | From 1 (very susceptible) to 9 (totally resistant) |

**Supplemental Table S3**. Genotypes sequenced for *in silico* chromosome painting and fine mapping of *Rpv2*

| Name | Phenotype | Fine mapping step | Type | ENA accession number |
|---|---|---|---|---|
| Trayshed | Resistant | Control | Rpv2 donor | SRR6729333 |
| 1771P | Resistant | Control | Female parent BC4 | ERR13382343,ERR13382342,ERR13382341,ERR13382340 |
| Nebbiolo | Susceptible | Control | Male parent BC4 | ERR13382347,ERR13382346,ERR13382345,ERR13382344 |
| 1126R | Resistant | Control | BC4 - Non recombinant | ERR13382351,ERR13382350,ERR13382349,ERR13382348 |
| 1142R | Susceptible | Control | BC4 - Non recombinant | ERR13382357,ERR13382356,ERR13382355,ERR13382354,ERR13382353,ERR13382352 |
| 1212R | Resistant | 780 kb | BC4 - Recombinant | ERR13382361,ERR13382360,ERR13382359,ERR13382358 |
| 1137R | Susceptible | 780 kb | BC4 - Recombinant | ERR13382365,ERR13382364,ERR13382363,ERR13382362 |
| 2859Z | Resistant | 780 kb | BC5 - Recombinant | ERR13382366 |
| 2862Z | Susceptible | 780 kb | BC5 - Recombinant | ERR13382367 |
| 2951Z | Susceptible | 780 kb | BC5 - Recombinant | ERR13382368 |
| 2952Z | Susceptible | 780 kb | BC5 - Recombinant | ERR13382369 |
| 2434D | Resistant | 250 kb | S - Recombinant | ERR13382370 |
| 3370D | Resistant | 250 kb | S - Recombinant | ERR13382371 |
| 3561D | Resistant | 250 kb | S - Recombinant | ERR13382372 |
| 4041D | Resistant | 250 kb | S - Recombinant | ERR13382373 |
| 2935D | Susceptible | 250 kb | S - Recombinant | ERR13382374 |
| 3798D | Susceptible | 250 kb | S - Recombinant | ERR13382375 |
| 4083D | Susceptible | 250 kb | S - Recombinant | ERR13382376 |

**Supplemental Table S4**. QTLs for resistance to downy mildew detected in the pseudo-BC1 population.

| Trait | LG | LOD | % Expl | Threshold | Peak (cM) | -1lod U (cM) | -1lod D (cM) | CI (cM) |
|---|---|---|---|---|---|---|---|---|
| **OIV452** | 12 | 4.32 | 12.6 | 1.6 | 56.73 | 50.13 | 74 | 23.87 |
| **OIV452** | 15 | 1.74 | 5.4 | 1.5 | 25.81 | 20 | 27.5 | 7.5 |
| **OIV452** | 18 | 13.98 | 36.4 | 1.6 | 61.76 | 57 | 67.4 | 10.4 |
| **S** | 18 | 22.73 | 55.6 | 1.6 | 61.26 | 58 | 63 | 5 |
| **SQ** | 7 | 2.75 | 20.5 | 1.4 | 6 | 0 | 25 | 25 |
| | 12 | 8.49 | 41 | 1.5 | 57 | 53 | 66 | 13 |
| | 15 | 1.71 | 9.7 | 1.4 | 26 | 19.5 | 27.4 | 7.9 |
| **DS** | 12 | 2.88 | 9.7 | 2.5 | 63 | 53 | 77 | 24 |
| | 18 | 17.78 | 45.9 | 2.7 | 61.2 | 59 | 64 | 5 |
| **NSTO** | 13 | 3.35 | 14 | 2.4 | 11 | 0 | 23.5 | 23.5 |
| | 18 | 9.44 | 26.8 | 2.7 | 64.4 | 59.3 | 69 | 9.7 |
| **NBSCP** | 12 | 6.19 | 17.2 | 2.55 | 56.7 | 53 | 66 | 13 |
| | 18 | 5.04 | 15.3 | 2.6 | 60.2 | 52 | 70 | 18 |

**LG**: linkage group; **% Expl**: percentage of the variance explained by the QTL; **Threshold**: significance threshold for QTL detection based on permutation test; **Peak**: position of the QTL peak; **-1lod**: position with a LOD value of (LODmax-1), upstream (**U**) and downstream (**D**) of the QTL peak. **CI**: confidence interval size.

**Supplemental Table S5**. Newly developed microsatellite markers in the *Rpv2* genomic region.

| Marker | Primer name | Primer sequence |
|---|---|---|
| Chr18.2000 | Chr18.2000 Fw | GGCTAGAACAAGGGTAGAAATG |
| | Chr18.2000 Rv | CAGATCGAAGAACGAAATTAGG |
| Chr18.2252 | Chr18.2252 Fw | TCTATCTACTTCTGTTTTGGGG |
| | Chr18.2252 Rv | AACCTCTCCCAGTAAAATCTTC |
| Chr18.2324 | Chr18.2324 Fw | ACTATGTTACTGTCCCCATTGA |
| | Chr18.2324 Rv | GGTTGACGTATGATGATTGAAG |
| Chr18.2336 | Chr18.2336 Fw | TGTTTCTGGTTTGAAGGTCTAC |
| | Chr18.2336 Rv | ACCATTAGGAAGACAGATAGGC |

**Supplemental Table S6.** Metrics of the assembly of the genome of *V. rotundifolia* cv Regale.

| | |
|---|---|
| **Number of contigs** | 1,572 |
| **Contig minimal size (bp)** | 19,372 |
| **Contig average size (bp)** | 274,669 |
| **Contig median size (bp)** | 103,511 |
| **N50** | 688,346 |
| **Contig maximal size (bp)** | 5,300,691 |
| **Size of the assembly (bp)** | 431,780,342 |
| **% complete genes based on BUSCO analysis** | 96.6% |

**Supplemental Table S7**. Genes present in the 250 kb interval containing *Rpv2*.

| Name | Fonction | Domains | CDS length (bp) | Position on chr18 (nt) |
|---|---|---|---|---|
| **231950.1.2** | Sulfotransferase | Sulfotransferase | 990 | 22425769-22426758 |
| **231950.3** | Cytochrome P450 | Plant_Cytochrome_P450_Monooxygenase | 645 | 22435391-22437694 |
| **231951** | NLR | NB-LRR | 1518 | 22451794-22470598 |
| **231960** | Cytochrome oxydase | Cytochrome C oxydase | 456 | 22471126-22472116 |
| **231970** | NLR | TIR-NBS-LRR | 3624 | 22484537-22491415 |
| **231981** | NLR | TIR-NB | 1360 | 22503743-22507089 |
| **231982** | NLR | LRR | 816 | 22511562-22525765 |
| **231990** | NLR | LRR | 1998 | 22545290-22589908 |
| **232010** | NLR | TIR-NBS-LRR | 3807 | 22607942-22613726 |
| **232011** | NLR | TIR | 393 | 22619425-22623301 |
